# Megalin (LRP2), prenatal betamethasone, and injury susceptibility in the developing kidney

**DOI:** 10.64898/2026.06.02.729064

**Authors:** Cimran Nakum, Brandi Bull, Sunitha Yarlagadda, Shalini Indugula, Kaitlynn Stowers, Katherine VandenHeuvel, Jenna T Ference-Salo, Jeffrey A Beamish, Avery Volz, J Elliott Robinson, Aisaku Nakamura, Lili Ding, Dilip K Singh, Bhagwat Prasad, Meredith P Schuh

## Abstract

Preterm infants undergo postnatal nephrogenesis and are often exposed to gentamicin (gent). Mothers at risk of preterm birth receive betamethasone (beta) to accelerate fetal lung development. Gent cytotoxicity occurs in proximal tubules (PT) after megalin (LRP2)-mediated endocytosis and accumulation. The objective of this study was to evaluate the impact of proximal tubular maturation, impacted by both age and prenatal beta, on injury susceptibility and nephron number in rats. Pups were given toxic gent dosing (100mg/kg) or saline intraperitoneal for 5 days during nephrogenesis (P0-4) or tubular maturation (P6-10). This was repeated with maternal exposure to beta to evaluate impact of beta on injury. Quantitative proteomics analysis revealed significantly higher levels of LRP2 at P10 as compared to P4, correlating with increased injury to gent exposure from P6-10 relative to P0-4. P10 pups exposed to prenatal beta had significantly more LRP2 protein relative to controls, which correlated to more injury after gent exposure at P6-10. Only those exposed to prenatal beta with P6-10 gent demonstrated ∼50% nephron reduction. This study supports that tubular maturation is a critical period of vulnerability to gentamicin correlating to LRP2 protein abundance. Prenatal corticosteroids increase the severity of acute and chronic injury in this highest risk exposure group.

## Introduction

Human nephrogenesis ends prior to birth in full-term infants, 34-36 weeks of gestation (GW),(1) with most nephrons (∼60%) being formed during late gestation, however, the preterm infant must continue this process postnatally,(2–4) which renders the immature kidney vulnerable to pharmacological insult. Preterm birth is associated with premature cessation of nephrogenesis, low nephron endowment, and an increased risk of chronic kidney disease (CKD) later in life.(2, 5–8) Preterm infants undergoing postnatal nephrogenesis are exposed to drugs that may impact both kidney development and injury risk.(8–10) Over 90% of all neonates under 28 weeks gestation are exposed to nephrotoxic medications,(11) with gentamicin (gent), used to treat neonatal sepsis, being the most common.(12) Furthermore, maternal treatment with prenatal steroids is associated with accelerated glomerular and kidney tubular maturation,(13–16) which could alter susceptibility based on this maturation. In a rabbit model, we found that nephrotoxin exposure at one week of life (including gentamicin) resulted in greater fibrosis and evidence of CKD at 3 months of life than exposure immediately after birth, suggesting increased vulnerability during tubular maturation and drug transporter development.(17) Megalin (LRP2) is a ∼600kDa glycoprotein belonging to the LDL receptor family which is found on the apical membrane of the proximal tubule epithelial cells.(18) In addition to its role in reabsorption of filtered proteins,(19) it also mediates the uptake of several nephrotoxic drugs, including gentamicin, vancomycin, colistin, and cisplatin.(20) During kidney development, LRP2 is one of the earliest transport proteins to be expressed and appears prior to other markers of tubular maturation.(21–23) Gentamicin cytotoxicity occurs after LRP2-mediated endocytosis into the proximal tubular cells, leading to intracellular accumulation of the drug.(24) Following uptake, gentamicin is trafficked and accumulates within the lysosomes(25) where lysosomal rupture contributes to inflammation, oxidative stress, and cell death, due to the proximal tubules lacking an efficient mechanism to get gentamicin out of the cells.(26–33) Since gentamicin has a low serum protein binding rate,(34) much of the circulating drug remains in free form and is filtered to the LRP2+ proximal tubule cells. In humans, LRP2 expression increases from 24 weeks gestation to full term birth.(21) Kidney tubular maturation occurs in a centrifugal pattern during development, with the inner cortex maturing earlier than the outer cortex, paralleling progressive increase in transporter expression, including LRP2.

In addition to the nephrotoxic gent exposures within the neonatal intensive care unit (NICU), preterm infants are often exposed to corticosteroids, such as betamethasone (beta) via prenatal exposure. Women at risk for preterm delivery receive prenatal beta to accelerate fetal lung maturation and improve neonatal survival, however, it coincides with the peak burst of in utero nephrogenesis during late gestation.(35–37) Clinical and translational work suggests that corticosteroids may negatively alter kidney development, influencing nephron development, and tubular differentiation/structure leading to long-term kidney harm.(38, 39)

In the present study, we utilized a rat model to evaluate the stage-specific effects of postnatal gent exposures on injury, recovery, and long-term outcomes. The rat demonstrates a prolonged window of postnatal nephrogenesis similar to the preterm infant, and undergoes a period of arcading nephrogenesis, a form of human nephron formation in extremely preterm infants, supporting its translatability to humans.(40) Specifically, nephrogenesis continues until postnatal day 5 (P5) and tubular maturation until P10 (**Supplemental Fig. 1**). Therefore, this model allowed us to test two nephron developmental periods in humans. P0-4 (<1 week of life) in the rat represents nephrogenesis at ∼24-26 weeks gestation (wg) in a preterm neonate, while P6-10 (∼1 week of life), represents tubular maturation at ∼32-36 wg in a preterm neonate. Here, we demonstrate how developmental timing of nephrotoxin exposure and prenatal corticosteroid administration interact to shape the trajectory of acute kidney injury (AKI) to CKD in the neonate, including changes in nephron number and fibrosis. We found that LRP2 protein abundance increases with age during early postnatal life but decreases in adulthood, and that prenatal corticosteroids administered to the mother perturb LRP2 levels. Furthermore, those exposed to corticosteroids prenatally and gentamicin at one week of life had the highest levels of fibrosis at 3 months of age, suggesting a difference in injury susceptibility. This study has broader applicability to LRP2-mediated nephrotoxicity and how external variables, such as corticosteroids, impact vulnerability.

## Results

### Gentamicin exposure during tubular maturation is more nephrotoxic than exposure immediately after birth and correlates to LRP2 expression

To determine if the developmental timing of nephrotoxin exposure influences the severity of kidney injury, toxic dosing of gent (100mg/kg)(41) was given immediately after birth (during nephrogenesis; P0-4) compared to one week of life (tubular maturation; P6-10) (**Supplemental Fig. 2**). We found that exposure from P0-4 produced minimal injury to the developing proximal tubules (**Fig. 1 A,C, Supplemental Fig. 3A**), while the same 100mg/kg gent exposure at one week of life (P6-10) led to a severe injury phenotype at the cortico-medullary border, evidenced by proximal tubule epithelial cell-specific kidney injury marker 1 (KIM1)(42) immunofluorescent (IF) staining (**Fig. 1 B, D, D^1^, Supplemental Fig. 3B**). This injury pattern is distinct from what is seen in adult rats (4-6 months of age) who similarly received 100mg/kg gent daily for five days, where KIM1 is located in the outer cortex but not the cortico-medullary border (**Fig. 1F**). These findings suggest that susceptibility of the developing kidney to nephrotoxin injury increases as tubular maturation and drug transporter development progresses.(17) To further explore the differential protein abundance of LRP2 and associated toxicity in the neonatal period, we performed proteomic measurement of LRP2 on kidney samples from rats at P4 (nephrogenesis), P10 (tubular maturation), and adulthood in controls and those who received 5 days of toxic gent exposure. Proteomic analyses identified increased LRP2 protein abundance per cell at P10 relative to P4 and adulthood (6110 vs 3828 pmol/mg membrane protein, p=0.0005; Table 2, **Fig. 1E**), suggesting a heightened period of non-monotonic expression of LRP2 corresponding to the peak injury period.

**Figure 1:**
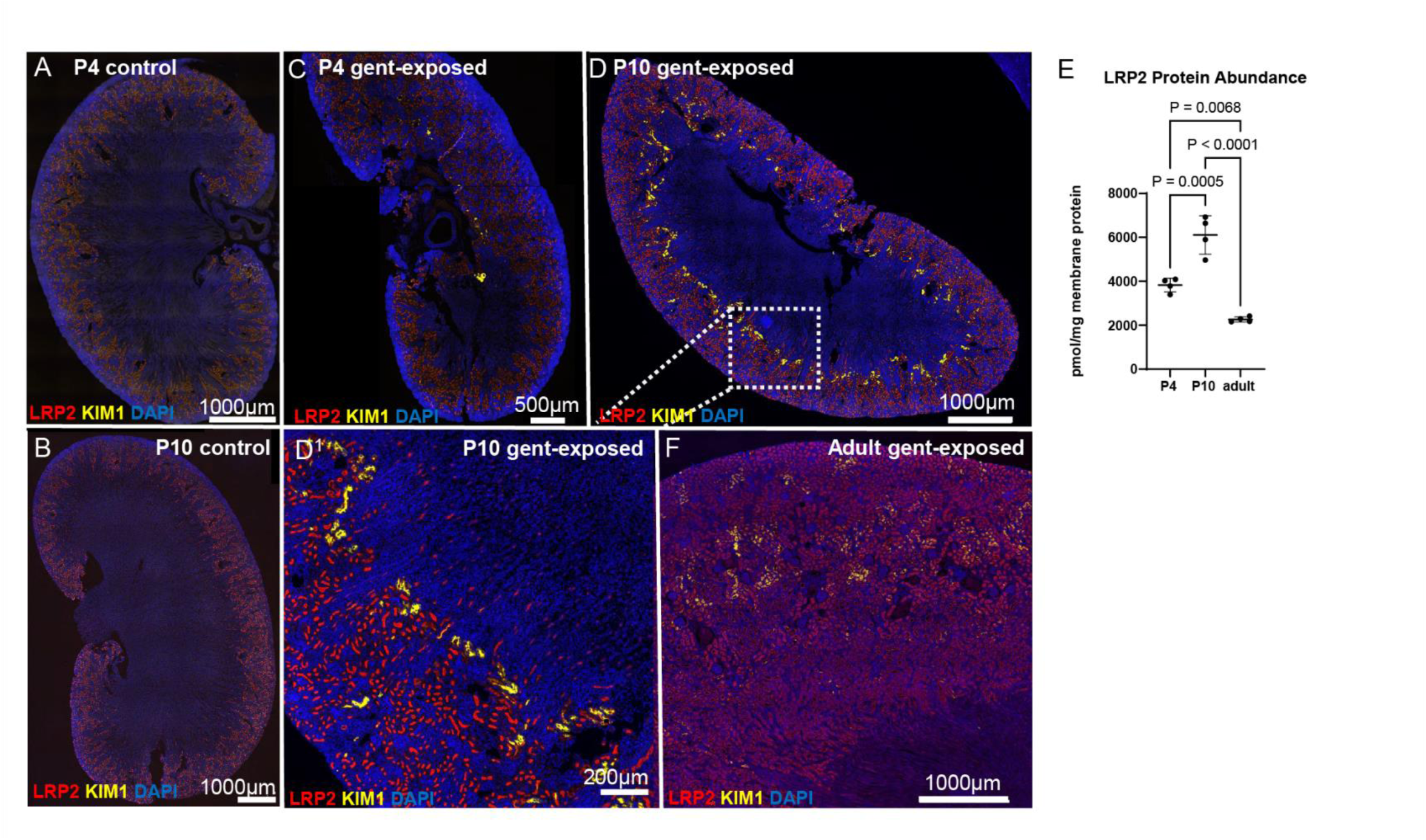
Nephrotoxic AKI worsens with increasing postnatal age but is distinct from adult kidney pattern of injury. **A)** P4 and **B)** P10 control kidney. **C)** Minimal KIM1 (yellow) staining seen in P4 **D)** Extensive injury is evident at P10, as demonstrated by KIM1 staining. Injury is located at the cortico-medullary border (D**^1^**) and co-stains with LRP2 (red). **E)** LRP2 protein abundance per cell is significantly higher at P10 compared to P4 or adult rats (n = 4 per group). **F)** This injury pattern is distinct from adult pattern of injury after gentamicin.

**Table 1:** Nephron Quantification (light sheet)

| Prenatal treatment | Postnatal treatment | Treatment period | Sample | Total number of gloms |
| --- | --- | --- | --- | --- |
| None | Saline | P0-4 | R11025 2C | 22648 |
| None | Saline | P0-4 | R11025 4C | 40312 |
| None | Saline | P0-4 | R11025 6C | 41109 |
| None | Saline | P0-4 | R11025 8C | 24535 |
| None | Gent | P0-4 | R11025 1E | 27111 |
| None | Gent | P0-4 | R11025 5E | 22365 |
| None | Gent | P0-4 | R11025 7E | 24487 |
| None | Gent | P0-4 | R11025 9E | 34951 |
| Beta | Saline | P0-4 | TS21825 2C | 23557 |
| Beta | Saline | P0-4 | TS21825 10C | 34437 |
| Beta | Saline | P0-4 | WS31925 6C | 31402 |
| Beta | Saline | P0-4 | WS31925 10C | 37007 |
| Beta | Gent | P0-4 | TS21825 1E | 29018 |
| Beta | Gent | P0-4 | TS21825 7E | 23023 |
| Beta | Gent | P0-4 | TS21825 13E | 33179 |
| None | Saline | P6-10 | F91224 6C | 35891 |
| None | Saline | P6-10 | F91224 10C | 47404 |
| None | Saline | P6-10 | F91224 14C | 43446 |
| None | Gent | P6-10 | F91224 3E | 18427 |
| None | Gent | P6-10 | F91224 5E | 36291 |
| None | Gent | P6-10 | F91224 15E | 20298 |
| Beta | Saline | P6-10 | SS20525 6C | 27507 |
| Beta | Gent | P6-10 | SS20525 3E | 8756 |
| Beta | Gent | P6-10 | SS20525 7E | 17297 |
| Beta | Gent | P6-10 | SS20525 9E | 11796 |
| Beta | Gent | P6-10 | SS20525 11E | 12345 |

**Table 2.** Summary Statistics for Outcomes.

| Figure | Variable | Statistics | Categories |  |  |  |  |  | Overall P |
| --- | --- | --- | --- | --- | --- | --- | --- | --- | --- |
| Figure 1E | LRP2 Protein Abundance (pmol/mg) | Categories (n) | P0-4 (4) | P6-10 (4) | Adult(4) | NA | NA | NA | <0.0001 |
|  |  | Mean(SD) | 3828 (314.3) | 6110 (872.0) | 2265 (117.2) | NA | NA | NA |  |
| Figure 2A | KW/BW (mg/g) | Categories (n) | Control(6) | P0-4(5) | NA | NA | NA | NA | 0.838 |
|  |  | Mean(SD) | 6.503 (0.7093) | 6.625 (1.113) | NA | NA | NA | NA |  |
| Figure 2B | KW/BW (mg/g) | Categories (n) | Control(7) | P6-10(6) | NA | NA | NA | NA | <0.0001 |
|  |  | Mean(SD) | 6.989 (0.8173) | 10.54 (1.110) | NA | NA | NA | NA |  |
| Figure 3 A | Blood Urea Nitrogen (mg/dL) | Categories (n) | Control P0-4(5) | Control P6-10(5) | Gent P0-4(5) | Gent P6-10(4) | NA | NA | 0.1339 |
|  |  | Mean (SD) | 19.4 (2.302) | 18 (1.000) | 20.8 (1.483) | 18.25 (2.630) | NA | NA |  |
| Figure 3B | Plasma Creatinine (mg/dL) | Categories (n) | Control P0-4(5) | Control P6-10(5) | Gent P0-4(5) | Gent P6-10(4) | NA | NA | 0.5867 |
|  |  | Median(IQR) | 0.3 (0.25,0.4) | 0.3 (0.25,0.45) | 0.4 (0.35,0.4) | 0.35 (0.3,0.475) | NA | NA |  |
| Figure 3H | Glomerular Count | Categories (n) | Control P0-4(4) | Control P6-10(3) | Gent P0-4(3) | Gent P6-10(3) | NA | NA | 0.1306 |
|  |  | Mean(SD) | 32151 (9919) | 42247 (5849) | 27268 (6738) | 25005 (9818) | NA | NA |  |
| Figure 4A | BUN (mg/dL) | Categories (n) | Control (5) | Gent P6-10(4) | Beta(4) | Beta+Gent P6-10(4) | NA | NA | <0.0001 |
|  |  | Median(IQR) | 18 (17,19) | 17.5 (16.25,21) | 23 (20.75,23) | 37.5 (34.5,48) | NA | NA |  |
| Figure 4B | Plasma Creatinine (mg/dL) | Categories (n) | Control (5) | Gent P6-10(4) | Beta(4) | Beta+Gent P6-10(4) | NA | NA | 0.0449 |
|  |  | Median(IQR) | 0.3 (0.25,0.45) | 0.35 (0.3,0.475) | 0.45 (0.4,0.5) | 0.6 (0.45,0.825) | NA | NA |  |
| Figure 4C | Nephron Number at age 3 (count) | Categories (n) | Control(7) | Gent P0-4(4) | Gent P6-10(3) | Beta(5) | Beta+Gent P0-4(3) | Beta+Gent P6-10(4) | 0.0016 |
|  |  | Mean(SD) | 36478 (9472) | 27229 (5502) | 25005 (9818) | 30782 (5372) | 28407 (5106) | 12549 (3537) |  |
| Figure 4D | Median glomerular volume per kidney ($\mu\text{m}^3$) | Categories (n) | Control(7) | Gent P0-4(4) | Gent P6-10(3) | Beta(5) | Beta+Gent P0-4(3) | Beta+Gent P6-10(4) | 0.0119 |
|  |  | Median(IQR) | 1537719 (1412319, 1729929) | 1478037 (1429542, 1575154) | 2115613 (1930193, 2116718) | 1557085 (1451586, 2093997) | 1547137 (1443797, 1728815) | 2561761 (2249584, 2743623) |  |
| Figure 4E | Glomerular density (1/mm<sup>2</sup>) | Categories (n) | Control(8) | Gent P0-4(5) | Gent P6-10(4) | Beta(10) | Beta+Gent P0-4(6) | Beta+Gent P6-10(4) | 0.0066 |
|  |  | Median(IQR) | 4.600 (4.061,6.048) | 5.875 (5.395, 7.565) | 5.327 (3.782, 6.086) | 4.305 (3.546, 5.306) | 4.684 (4.173, 5.187) | 2.507 (2.493, 3.382) |  |
| Figure 5A | LRP2 Normalized to β-Actin | Categories (n) | P10 Control(5) | P10 Beta(5) | NA | NA | NA | NA | 0.0317 |
|  |  | Median(IQR) | 0.1803 (0.1489, 0.4331) | 0.6161 (0.4143, 0.6555) | NA | NA | NA | NA |  |
| Figure 5D | Kim1 fold change | Categories (n) | Gent P6-10(7) | Beta+Gent P6-10(7) | NA | NA | NA | NA | 0.3829* |
|  |  | Mean(SD) | 16.92 (10.52) | 61.76 (51.97) | NA | NA | NA | NA |  |
| Figure 5E | Apoptotic Nuclei | Categories (n) | Control (4) | Gent P6-10(4) | Beta(4) | Beta+Gent P6-10(4) | NA | NA | 0.01 |
|  |  | Mean(SD) | 17.56 (9.846) | 73.37 (27.66) | 30.27 (2.025) | 96.38 (52.67) | NA | NA |  |

\*The P-value for Figure 5 D was calculated comparing delta Ct.

### Ongoing injury present two weeks after neonatal nephrotoxic exposure

To assess the impact of injury timing on structural response following AKI, we measured kidney weight normalized to body weight (KW:BW) two weeks after last gent exposure during early or late postnatal exposure windows. Consistent with the limited injury observed after early exposure, the P0-4 treated rats exhibit KW:BW ratios that are not significantly different to control treated pups (p=0.838; Table 2, **Fig. 2A**). In contrast, gent exposure at the late stage during tubular maturation resulted in a significant increase in KW:BW compared to control pups (10.54 vs 6.989 mg/g, p<0.0001; Table 2, **Fig. 2B**). To characterize the cellular response and ongoing injury following gent exposure, kidneys were examined via IF staining two weeks after initial injury. KIM1 staining was intermittently seen in P0-4 treated rats near the corticomedullary border, co-localizing with LRP2 (**Fig. 2C, C^1^**). More extensive damage was seen throughout the cortex of the gent P6-10 exposed pups, with evidence of tubular dilatation and diffuse KIM1 staining. (**Fig. 2D, D^1^**). We next investigated transcription factor SOX9 expression in these two-week samples. SOX9 is associated with tubular dedifferentiation and epithelial stress.(43–45) After injury, SOX9 is induced to help restore cell-cell contacts and apical-basal polarity. We identified increased nuclear SOX9 expression throughout the late-exposed proximal tubules compared to early-exposed (**Fig. 2C^1^, D^1^**). Adult rats treated with gent for 5 days had minimal injury at 2 weeks post-injury, with sparse KIM1 being seen via IF when compared to adult controls (**Fig. 2E^1^, F^1^**), suggesting prolonged recovery in the neonatal rats compared to adults.

**Figure 2:**
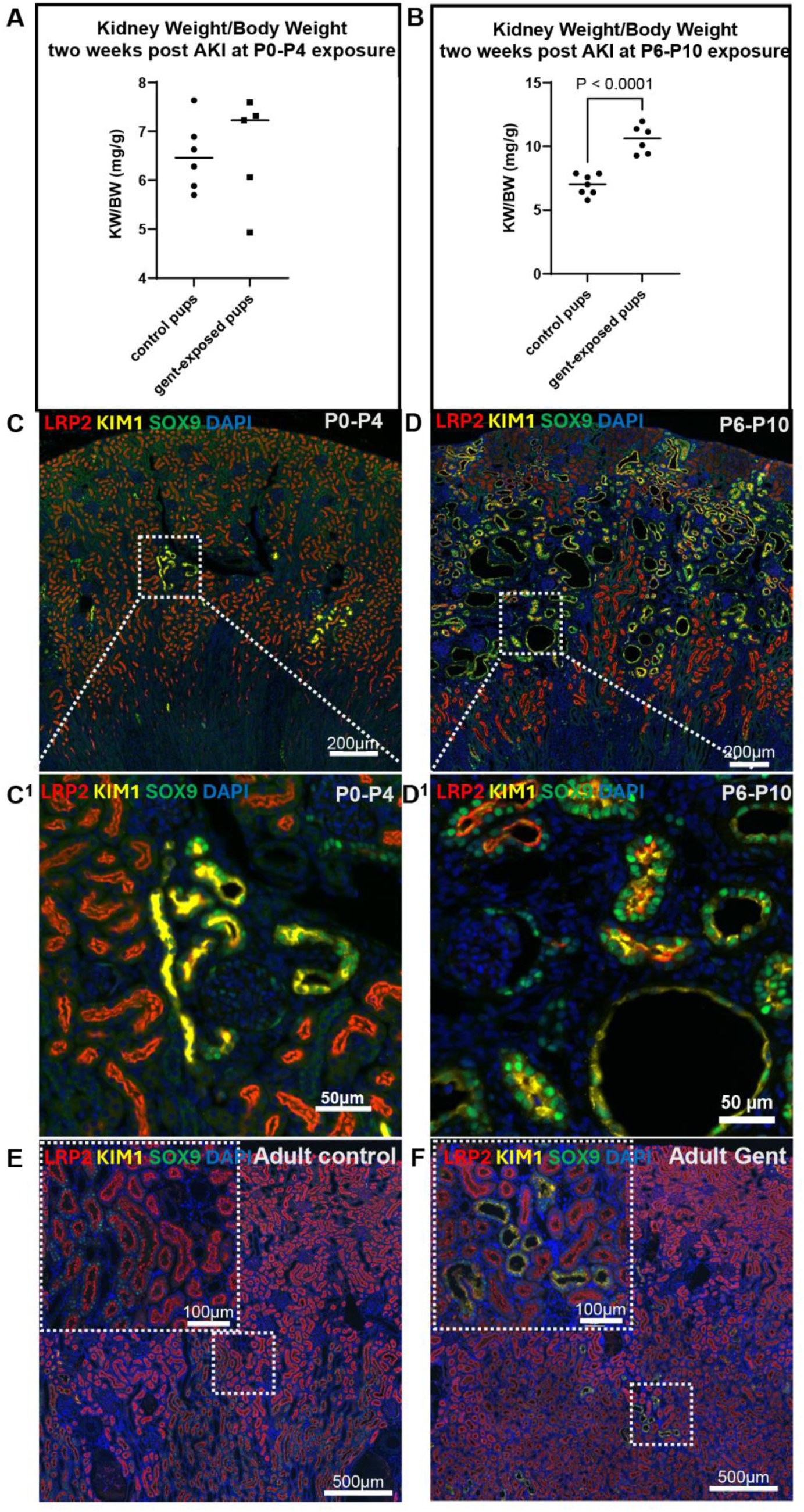
P10 animals demonstrate ongoing injury two weeks after toxic gentamicin exposure. Two weeks post gent exposure, kidney weight to body weight was assessed. **A)** no statistically significant difference was seen in the gent P0-4 group compared to controls (p=0.829) but **B)** gent P6-10 kidneys were significantly larger than controls. **C-C^1^)** Minimal damage was identified in gent P0-4 kidneys, but **D-D^1^)** extensive tubular damage with KIM1+SOX9+ staining was seen in gent P6-10 kidneys. **E-F)** this injury pattern is distinct from the adult rat.

### Nephron number and biochemical measurements of kidney function are stable at 3 months of age

Blood urea nitrogen (BUN) and plasma creatinine (PCr) were measured at three months of age. Despite the severe injury seen at two weeks, there was minimal biochemical evidence of injury at 3 months of age. The average BUN was 20.8mg/dL in the gent P0-4 rats and 18.3 mg/dL in the gent P6-10 rats, not significantly different from controls (overall p=0.1339; Table 2, **Fig. 3A**). PCr levels did not significantly differ among all four groups (overall p=0.5867; Table 2, **Fig. 3B**). We assessed fibrosis using Masson’s Trichrome staining and found that all groups demonstrated minimal to no fibrosis (**Fig. 3C-F**). Using Tomato Lectin perfusion, tissue clearing, and imaging using light sheet microscopy,(46) (**Supplemental Movie 1**) we quantified nephron number based on perfusion to the glomerular basement membrane from tomato lectin. Nephron numbers did not significantly differ among all groups, supporting resilience and recovery despite initial results (overall p=0.1306; Table 2, **Fig. 3H**).

**Figure 3:**
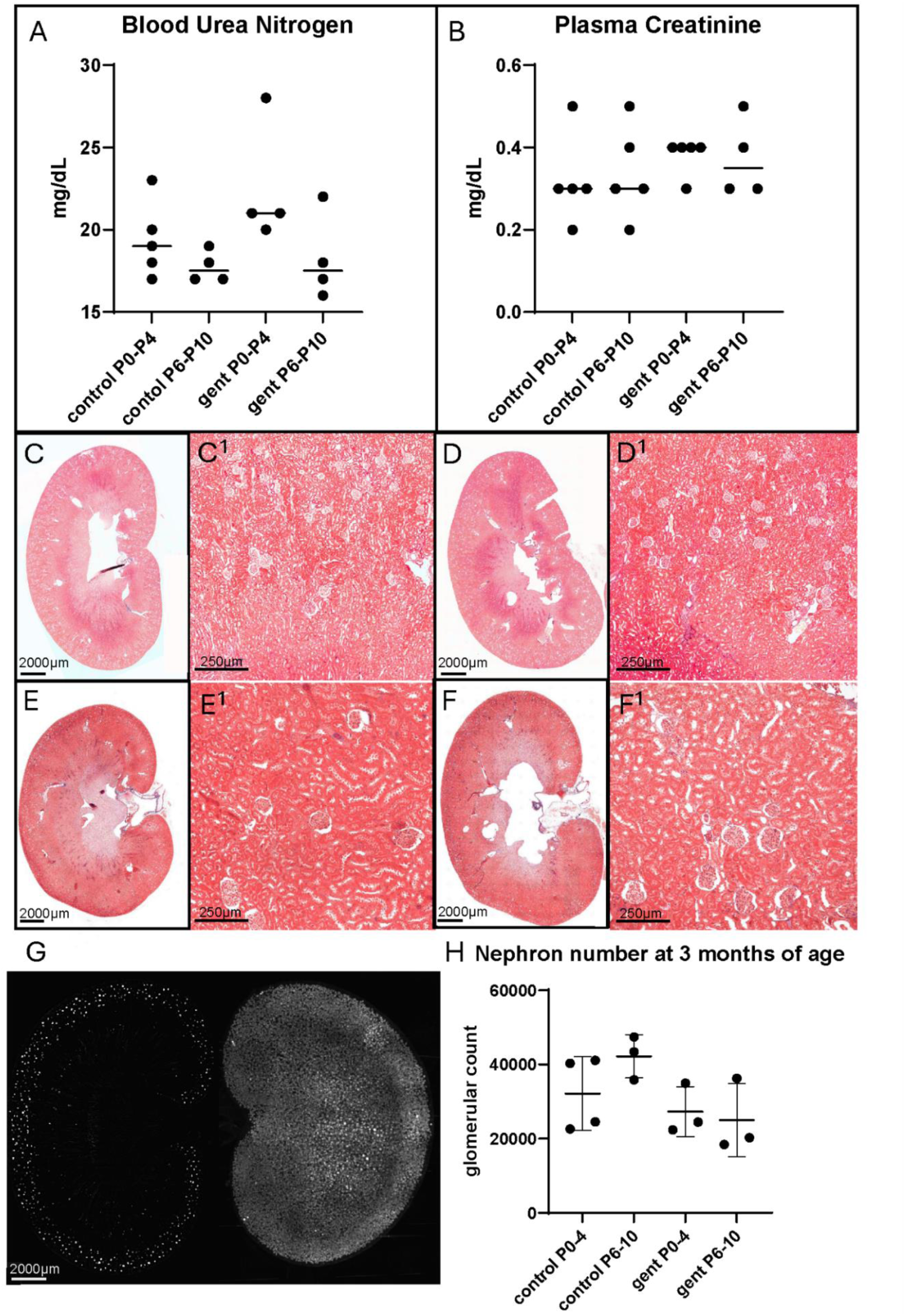
Animals recover after toxic gentamicin exposure. A-B) No significant difference was identified in BUN and Plasma creatinine levels in either exposure group. **C-F)** Minimal fibrosis was identified with similar findings between all groups, further supporting recovery. **G)** Example of light sheet microscopy quantification of nephron number after clearing **H)**. Glomerular counts using light sheet. Trend towards lower nephron counts after gent in both groups but still similar to controls.

In summary, toxic gent exposure during tubular maturation causes more injury than during nephrogenesis, and the pattern of injury is distinct from the adult. Despite extensive injury at two weeks after exposure, kidneys in both exposure timing groups demonstrated recovery at 3 months of age, with relatively stable BUN, PCr, and nephron number between all groups. We next sought to identify how prenatal beta exposure, a near universal exposure in infants born preterm, alters this window of susceptibility to injury and long-term kidney health.

### Does prenatal betamethasone alter susceptibility to gentamicin nephrotoxicity?

To determine whether prenatal beta altered susceptibility to injury and long-term functional impairment, we first performed a top-down approach, assessing 3-month outcomes. Now, pregnant dams were exposed to betamethasone at embryonic (E) day 17 and 18, with the gentamicin exposure periods replicated as above (**Supplemental Fig. 2**).

### Prenatal betamethasone and early (P0-4) gentamicin exposure: No difference in long-term outcomes

BUN and PCr at three months of age were compared between saline controls, gent P0-4 exposure, prenatal beta only, and prenatal beta with gent P0-4 gent. BUN, PCr, and nephron number were comparable across all four groups (**Supplemental Fig. 4 A-B**). There was a trend of increased BUN in prenatal beta (23mg/dL) and beta + gent P0-4 (23.17mg/dL), relative to control (19.4mg/dL) or gent P0-4 (20.8mg/dL). However, no significant difference was identified after multiple comparisons. Similarly, fibrosis was assessed which was similar between all four groups (**Supplemental Fig. 4 D, E**). Nephron number was compared, and there was nearly no difference in nephron endowment between groups and the nephron number was comparable to a healthy/normal adult rat (p = 0.719, **Supplemental Fig. 4 C**).(47)

### Prenatal betamethasone and late (P6-10) gentamicin exposure: Kidney dysfunction, fibrosis, and low nephron number at three months of age

Similar to the P0-4 long-term outcomes above, kidney structure and function were assessed in adulthood (3 months of age) following prenatal beta exposure and postnatal gent P6-10 exposure. Unlike all previous comparisons, animals exposed to prenatal beta + gent P6-10 displayed elevated BUN levels. Median BUN in beta + gent P6-10 was 37.5mg/dL compared to 18mg/dL (p=0.0147), 17.5mg/dL (p=0.0165), and 23mg/dL (p>0.99) in control, gent P6-10, and beta only respectively (Table 2, **Fig. 4A**). Notably there was still a trend of increased BUN in beta only group. No significant difference was seen between P6-10 treatment group PCr after multiple comparisons but trended towards increased PCr (0.6mg/dL compared to 0.3-0.45mg/dL) (**Fig. 4B**).

**Figure 4:**
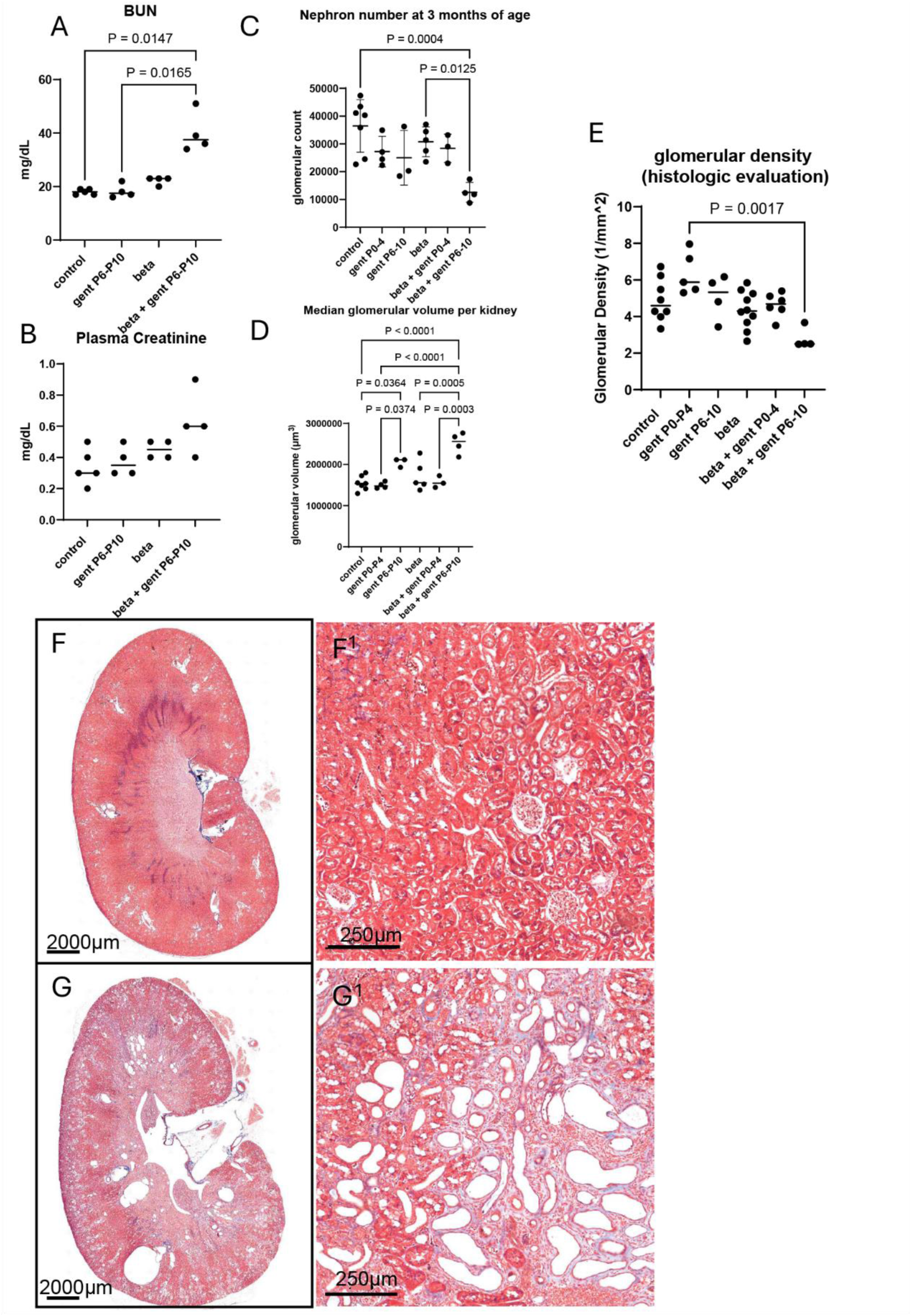
Prenatal betamethasone and Gent P6-10 result in long-term injury, fibrosis, low nephron number, and glomerular hypertrophy at 3 months of life. **A)** BUN significantly elevated in beta + gent P6-10 at three months of age. **B)** Trend in increased PCr in beta + gent P6-10 but not significant after multiple comparisons. **C-D)** Nephron number was 50% lower in beta + P6-10 kidneys, with associated demonstration of increased glomerular size compared to controls concerning for compensatory hypertrophy. **E)** Glomerular density was lower in beta + gent P6-10. **F-F^1^)** compared to beta alone, **G-G^1^)** beta + gent P6-10 demonstrate interstitial fibrosis, tubular dilatation, and scarring.

Histological analyses using Masson’s Trichrome staining demonstrated interstitial fibrosis in the beta + gent P6-10 group, with widespread structural abnormalities, including tubular dilatation, and increased interstitial fibrosis (**Fig. 4G, G^1^**). Kidneys from control and beta only groups displayed normal kidney architecture with minimal interstitial staining (**Fig. 4F, F^1^**). Kidneys underwent a semiquantitative scoring system by a clinical pathologist, which revealed that the beta + P6-10 group had a more widespread form of fibrosis, with most samples having severe fibrosis covering >50% of the cortex (**Supplemental Table 1)**. Nephron quantification at 3 months of age (n = 26) identified a statistically significant difference in nephron number (p = 0.0016, **Table 2**) between exposure groups. This significant reduction in nephron number was seen in the beta + P6-10 gent group (mean 12,549 ± 3,537 nephrons) compared to the control group (36, 478 ± 9472 nephrons, p = 0.0004) and compared to the prenatal beta alone group (30,782 ± 5,372 nephrons, = 0.0125 **Table 2**, **Fig.4 C**). Beta + P6-10 gent group also showed a substantial reduction (∼50%) compared with rats that received P6-10 gent alone (25,005 ± 9,818 nephrons), although this difference was not statistically significant (p = 0.2491). To verify our light sheet quantification, we measured glomerular density (n = 37) in parallel using previously established automated quantification method,(48) which was modified for the rat trichrome sections. Additionally, this allowed for quantification of 11 samples insufficiently cleared for light sheet quantification (**Supplemental Table 2**). Similarly, we identified a significant difference in glomerular density between groups (p=0.0066, **Table 2**). Glomerular density was decreased in the beta + P6-10 group (2.507) compared to gent P0-4 group (5.875, p=0.0017; Table 2, **Fig. 4E, Supplemental Table 2**). To evaluate functional compensation for low nephron number, we evaluated for difference in glomerular volume in the adult animals. We found significantly larger glomeruli in the beta + gent P6-10 animals (2,561,761 µm^3^) compared to controls (1,537,719 µm^3^, p<0.0001), compared to gent P0-4 (1,478,037 µm^3^, p < 0.0001), compared to beta alone (1,557,085 µm^3^, p = 0.0005), and compared to beta + gent P0-4 (1,547,137 µm^3^, p=0.0003), supporting compensatory hypertrophy due to low nephron number (**Table 2, Supplemental Table 3, Fig 4D**). Additionally, a significant difference was seen in gent P6-10 (2,115,613 µm^3^) compared to controls (p = 0.0364) and gent P0-4 (p = 0.0374).

### Increased LRP2 after beta as a potential mechanism for increased susceptibility to injury after P6-10 gentamicin exposure

To understand the reasons for our long-term findings, we next assessed whether prenatal betamethasone exposure alters 1) LRP2 abundance and 2) nephrotoxic injury from P6-10. Using the Western blot, quantification revealed a ∼ 2-fold increase in LRP2 protein (0.6161 vs 0.1803) in beta-exposed P10 kidneys compared to P10 controls (p = 0.0317), with representative immunoblots confirming increased protein abundance, all normalized to β-actin (Table 2, **Fig. 5 A, Supplemental Fig. 5**). These findings suggest that prenatal beta accelerates LRP2 abundance during the experimental exposure period.

**Figure 5:**
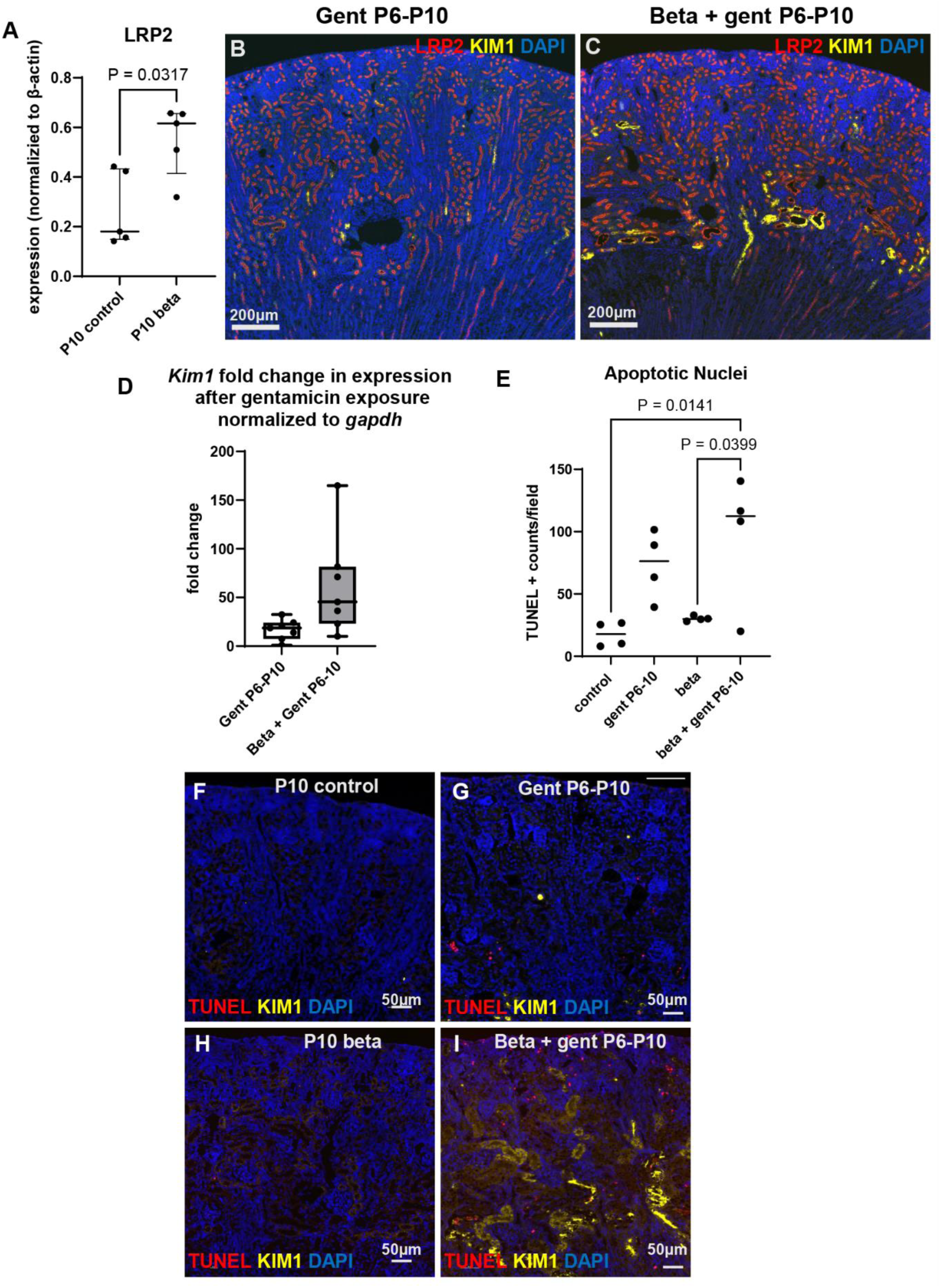
Prenatal beta increases LRP2 at P10, with increased injury and apoptosis after gentamicin exposure at P6-10. **A)**. LRP2 was significantly increased in P10 animals after prenatal beta compared to P10 controls on western blot analysis. **B-C)** we identified increased KIM1 staining at the cortico-medullary border of beta + gent P6-10 compared to gent P6-10. **D)** this finding was supported by increased fold change in *Kim1* RNA expression on RT-qPCR. **E)** Apoptotic nuclei were assessed using TUNEL assay. Representative staining showed in **F-I**.

We next compared AKI between gent P6-10 and beta + gent P6-10. While both demonstrate KIM1 staining at the corticomedullary border, there appeared to be more KIM1 staining in those that received prenatal beta (**Fig. 5 B-C, Supplemental Fig. 3C**). This was further analyzed using RT-qPCR, where we found higher fold change in *Kim1* expression in beta + gent P6-10 compared to P6-10 gent, relative to age matched controls. However, delta Ct values were not significantly different between these two groups (p=0.3829; Table 2, **Fig. 5 D**). We next evaluated differences in apoptosis as a cause for long-term injury and fibrosis using TUNEL assay. Increased apoptosis was seen in beta P6-10 (96.38 TUNEL+ counts/field) compared to controls (17.46 TUNEL+ counts/field, p = 0.0141) and compared to beta (30.27 TUNEL+ counts/field, p = 0.0399, Table 2, **Fig. 5 E**). There was a trend of increased BUN in gent P6-10 group compared to controls but this was not statistically significant (p = 0.091).

This increase in apoptosis seen in beta + gent P6-10 groups was also visible via the TUNEL stain as well and was accompanied by high amounts of KIM1 staining (**Fig. 5 F-I**). Together, these findings demonstrate that increased LRP2 abundance is seen after beta exposure and is also associated with increased KIM1 expression and apoptosis. While gent-induced nephrotoxic injury alone is largely recoverable, the combination of prenatal beta followed by toxic gent exposure during peak LRP2 abundance results in persistent kidney dysfunction and fibrosis in adulthood.

In the mixed effects model analysis, beta + gent P6-10 had significantly less nephrons compared to saline controls (estimated difference =-23578, adjusted p = 0.0068) and beta controls (estimated difference =-17376, adjusted p = 0.0354). Similarly, beta + gent P6-10 had significantly lower glomerular density when compared to saline controls (estimated difference =-2.1110, adjusted p= 0.013) (**Table 3**).

**Table 3:** Estimated Differences from the Linear Mixed-Effects Models.

| Response | Label | Estimated Difference | Standard Error | Lower 95% CI | Upper 95% CI | P-value | Bonferroni Adjusted P |
| --- | --- | --- | --- | --- | --- | --- | --- |
| Glomerular Count | None Saline vs Beta Gent P0-4 | 7824.05 | 6283.85 | -5433.72 | 21082 | 0.23 | 1 |
|  | None Saline vs Beta Gent P6-10 | 23578 | 6031.67 | 10852 | 36303 | 0.0011 | 0.0068 |
|  | Beta Gent P0-4 vs Beta Saline | -1622.25 | 5439.14 | -13098 | 9853.33 | 0.7691 | 1 |
|  | Beta Gent P6-10 vs Beta Saline | -17376 | 5524.28 | -29031 | -5720.68 | 0.0059 | 0.0354 |
|  | None Gent P0-4 vs None Saline | -7071.31 | 4546.98 | -16665 | 2521.99 | 0.1383 | 0.83 |
|  | None Gent P6-10 vs None Saline | -14377 | 5145.13 | -25232 | -3521.31 | 0.0125 | 0.0747 |
| Glomerular Density | None Saline vs Beta Gent P0-4 | 0.2737 | 0.5497 | -0.8541 | 1.4016 | 0.6226 | 1 |
|  | None Saline vs Beta Gent P6-10 | 2.111 | 0.6233 | 0.8321 | 3.3898 | 0.0022 | 0.0131 |
|  | Beta Gent P0-4 vs Beta Saline | 0.2724 | 0.5256 | -0.806 | 1.3508 | 0.6085 | 1 |
|  | Beta Gent P6-10 vs Beta Saline | -1.5649 | 0.6021 | -2.8004 | -0.3294 | 0.015 | 0.0898 |
|  | None Gent P0-4 vs None Saline | 1.4542 | 0.5802 | 0.2636 | 2.6447 | 0.0185 | 0.1112 |
|  | None Gent P6-10 vs None Saline | 0.1602 | 0.6233 | -1.1187 | 1.439 | 0.7991 | 1 |

## Discussion

In a neonatal rat model, we found that toxic (100 mg/kg) gentamicin exposure at one week of life (during proximal tubular maturation) led to a more severe injury phenotype than immediately after birth, correlating with increased LRP2 protein abundance per cell. Our most striking finding was that the combination of prenatal corticosteroids with postnatal gentamicin during tubular maturation (peak LRP2 abundance) led to the highest levels of fibrosis and biomarkers of CKD in adulthood compared with either exposure alone. This suggests that the combination and timing of exposures are associated with peak LRP2 abundance and the greatest toxicity, with long-term evidence of CKD.

In contrast to the gent P0-4 exposure, gent P6-10 treated rats developed a more pronounced injury that persisted two weeks after exposure. These findings are supported by increased LRP2 expression during this period. LRP2 is responsible for gentamicin entry into the cell, and therefore we hypothesized that higher LRP2 expression would result in more gent uptake into the neonatal kidney’s proximal tubular epithelial cells, leading to higher intracellular accumulation and contributing to a worse injury phenotype at P10. This enrichment within proximal tubules aligns with the known role of LRP2 in mediating aminoglycoside endocytosis and provides a structural correlation to the increased vulnerability to gent-induced injury at the P10 window of postnatal treatment.(49) This pattern supports a model in which maturation-dependent upregulation of LRP2-mediated endocytosis enhances intracellular accumulation of gentamicin, amplifying cellular stress and ongoing injury. In contrast to earlier developmental stages, where lower LRP2 abundance may limit drug intake, the P10 kidney is uniquely positioned for increased gentamicin uptake. We also demonstrate that gentamicin remains in the tissue two weeks after exposure in P10 animals, even after non-measurable plasma levels. LRP2 is a mediator of nephrotoxic drug uptake and accumulation within the proximal tubular cells, linking increased receptor activity to heightened susceptibility to kidney injury.(49, 50) Nephrotoxic medication exposure is highly prevalent in premature neonates and is associated with AKI development,(10, 51) but how this AKI timing is associated with stage of kidney development is not known. Preclinical models of low nephron number support increased gentamicin-induced nephrotoxicity(52) and may be related to altered maturation to compensate for the ex-utero environment.

### The role of prenatal steroids and vulnerability

We found that the combination of prenatal steroids at E17 & 18 with postnatal gent during tubular maturation led to increased injury at the early AKI analysis, as well as persistent injury, fibrosis, and nephron loss into adulthood, which was not seen in any of the other treatment groups. We identified an increase in LRP2 at the vulnerable P10 window after beta exposure compared to controls. The P10 kidney is uniquely positioned at a point where increased functional maturation leads to a higher vulnerability to nephrotoxin exposure, which is further heightened upon prenatal steroid exposure. Importantly, the increased injury is accompanied by a corresponding increase in tubular apoptosis upon gent exposure, as demonstrated by enhanced TUNEL+ nuclei.

By adulthood, the beta + P6-10 group demonstrated marked signs of kidney functional decline and reduced nephron number (∼50% loss) compared with all other treatment groups. This group also had near doubling in their BUN and diffuse fibrosis. In contrast, despite experiencing greater acute injury, the P10 gent-only group did not exhibit these long-term deficits, with preservation of nephron number, absence of fibrosis, and normalization of functional markers into adulthood. This divergence highlights that while the P10 kidney exists within a window of increased susceptibility to injury, this alone is insufficient to drive chronic kidney disease. Rather, the addition of prenatal corticosteroids appears to fundamentally alter the injury-repair balance. In this context, corticosteroid exposure likely amplifies the initial injury burden, potentially through increased LRP2 and proximal tubule maturation, which would enhance gentamicin uptake and apoptosis. Together, these findings support a model in which developmental timing establishes a vulnerable state, but it is the interaction with the perinatal environment that ultimately determines whether injury resolves or progresses to long-term structural and functional decline. Our results differ from Ortiz et al, who found a 20% reduction in nephron number from prenatal steroids (dexamethasone) alone at three months of life,(53) while our experiment required the “second hit” of gentamicin. Findings in both studies may be due to increased apoptotic signaling,(54) which may cause progression towards fibrosis and CKD outcomes.

This study has a broader significance with important implications for the clinical care given to both mother and the premature neonate. In current neonatal practice, prenatal corticosteroids are administered to a vast majority (>90%) of pregnancies that are at risk of preterm delivery, while gent is routinely used postnatally within the NICU for infection management.(12, 55, 56) This combination of prenatal steroid exposure and postnatal nephrotoxic insult modeled here reflects a highly common and real-world scenario. The interactions between beta and gent highlights how perinatal exposures can shape long-term kidney health and suggest that this combination may predispose an individual to dysfunctional repair in a clinically relevant patient population. In tandem with these observations, our findings identified a conserved window of increased vulnerability during tubular maturation, showing that the timing of exposure has a profound impact, not only on the severity of short-term injury, but also on the capacity of the neonatal kidney to repair and recover into adulthood. While injury occurs during both ongoing nephrogenesis and tubular maturation, the ability to recover from the acute injury differs between these developmental stages. Notably, with the widespread clinical use of steroids, our data implies that this non-recoverable phenotype may be more present in clinical practice than our model of recovery. Given that beta exposure alone did not impact nephron number, it is critical to re-evaluate the role of increased susceptibility, apoptosis, and nephron loss rather than early cessation as the cause of low nephron number in preterm infants.(2) This has important implications for neonatal care, as it suggests that exposure to nephrotoxic agents such as gent during tubular maturation, particularly in cases with prenatal steroid exposure, may shift AKI toward CKD. Additionally, a novel aspect of this study was the use of whole-kidney light sheet fluorescent microscopy, which was verified by established pipelines and allowed for three-dimensional assessment of functional nephrons and was further supported by histological findings of reduced glomerular density.

Our study has many limitations. Importantly, we could not identify sex of pups at P4, and this determination was performed later, which lead to unbalanced groups. However, we were able to achieve an n ≥ 3 males and females per experimental group, with most experimental groups having an n = 5 or more. Additionally, we did not perform measured glomerular filtration rates of the adult animals, so we do not know if these numbers reflect true decline in glomerular filtration rate (GFR). We also do not know the GFR of the neonatal animals at the time of exposure, and how the role of urinary flow in relation to GFR impacts LRP2-mediated endocytosis. In the rat model and in the human, the kidney forms in a centrifugal fashion, with more mature nephrons being at the cortical medullary border. Our timing of exposure during “ongoing nephrogenesis” and “tubular maturation” describe the nephrogenic zone but not the function or development of the deeper nephrons. Another limitation is that we provided toxic levels of gentamicin, and do not yet know if these findings would persist in more clinically relevant dosing. However, the toxic findings must be tested before variations in dosing are evaluated. Finally, we caution the clinical implication of these findings. These exposures in the neonatal period do not occur in a vacuum but rather are highly complex and depend on the clinical need of the patient. Gentamicin is first line for neonatal sepsis and should be used to improve overall survival.(12) Similarly, prenatal corticosteroids have significantly improved outcomes in preterm infants through reductions in mortality and respiratory morbidity, establishing them as a critical component of neonatal care.(57–59) However, as our neonatal survival rates continue to improve, it is important to reflect on our standards of care, especially if there is flexibility in timing or ability to monitor drug dosing. Future work will focus on non-invasive metrics of tubular maturation and drug monitoring to identify infants most at risk.

Overall, our findings demonstrated that embryonic beta exposure combined with toxic gent exposure during tubular maturation and peak LRP2 expression causes severe injury that progresses to persistent kidney damage in adulthood. This model highlights developmental timing as a key determinant of the AKI to CKD progression in drug-induced injury and provides new insight into how perinatal exposures impact that risk.

## Methods

### Animals

Timed-pregnant and bred pairing Sprague Dawley rats were used for this study, purchased from Charles-River Laboratories. They were maintained and taken care of as per IACUC protocols: 2022-0042 and 2025-0036, all methods were approved and followed as per the approved protocols and guidelines. All sample/rat IDs are listed in **Supplemental Table 4,** including sex identification. *Sex as a biological variable:* All rats were used regardless of sex. Our study examined male and female animals, and similar findings are reported for both sexes. Pups were randomly divided and numbered between the control and experiment groups for all analysis timepoints. Inclusion criteria for analysis included presence of two anatomically normal kidneys at time of collection. One pup (VC 9E) was excluded based on this criterion. Adult rats were euthanized by overdose of carbon dioxide exposure followed by lethal dose of pentobarbital and exsanguination. Pups used for the study were anesthetized using isoflurane and euthanized with an overdose of pentobarbital (65 mg/mL concentration, 0.1-0.5 mL depending on weight at collection time) and exsanguination.

Pregnant dams were anesthetized in a chamber and externally with a nose cone for prenatal intramuscular (IM) betamethasone treatments (Cat#NDC-0517-0720-01, 6 mg/mL stock concentration), with a dose of 0.25 mg/kg based on weights at time E17 and 18. Pups were treated from P0-4 or P6-10 with gentamicin via an intraperitoneal (IP) injection (Fresenius Kabi USA, Cat#6332301003, 40 mg/mL stock concentration diluted to a 10 mg/mL, with a dose of 100 mg/kg based on weights at time on injection), control pups received saline at the same dose and time. Betamethasone dosing was chosen based on previous literature and clinical correlation to human dosing.(53, 60, 61)

### Immunofluorescence

5µm sections of paraffin embedded tissues were used for immunofluorescence (IF) staining. IF was performed as previously described.(17) Primary antibodies used in this study included: DAPI (Cat#D1306, Invitrogen, 1:1000), Goat anti-Rat Kim1 (Cat#AF3689, R&D, 1:200), Rb Lrp2 (Cat#AB76969, Abcam, 1:100), Gp Krt8 (Cat#GPK-8, Progen, 1:500), Alexa Fluor647 conjugated Rb WT1 (Cat#AB202639, Abcam, 1:250), Shp CHD6 (Cat#AF2715, R&D, 1:500), and Rb Sox9 (Cat#NBP1-85551, Novus biologicals, 1:200). Secondary antibodies used in this study included: Donkey anti-Gt 750 (Cat#ab175745, Abcam, 1:250), Donkey anti-Rb 647 (Cat#711-606-152, Jackson ImmunoResearch, 1:250), Donkey anti-Ms 594 (Cat#715-586-151, Jackson ImmunoResearch, 1:250), and Donkey anti-Gp 488 (Cat#706-546-148, Jackson ImmunoResearch, 1:250). Images were acquired on the Widefield Nikon Ti2 Inverted Microscope with a 20x objective. Images were processed on NIS-Elements software, version AR 6.20.02.

### RT-qPCR

RT-qPCR was completed to assess Kim1 (Millipore Sigma) protein expression. One half of the right kidney was collected and kept in RNA Later (Cat#AM7020, Thermofisher Scientific) and stored at-20°C. Prior to RNA extraction, tissue from cortex was cut and homogenized in RNASTAT 60 (Cat#CS-110, AMSBio), and RNA was extracted using GeneJET RNA Purification Kit (K0732, Thermofisher Scientific). Protocol used combined AMSBio RNASTAT 60 and GeneJET steps. RNA was quantified using the Nanodrop 2000C (Thermoscientific). cDNA was synthesized using SuperScript™ IV First-Strand Synthesis System (Cat#18091050, Thermofisher Scientific), manufacturer protocol was used. qPCR was performed using Powerup SYBR green Mastermix (Cat#A25742, Thermofisher Scientific) in a QuantStudio 5 Realtime PCR system. Samples were normalized to *Gapdh* and *Ppia* (Millipore Sigma) and fold change was calculated between age-matched control pups and experiment pups and normalized to housekeeping genes, *Gapdh* and *Ppia*. Primer sequences found in **Supplemental Table 5.**

### Protein Extraction & Western Blot

One half of the right kidney was collected, snap frozen and stored at-80°C. Prior to protein extraction, one half of kidney was used for P4 and P10 collections given small size and to maintain similar extraction protocols. Two week tissue samples were first cut in half and then dissected to include entire depth of cortex. Tissue was lysed in buffer made of: RIPA Lysis Buffer (Cat#89900, Thermofisher Scientific) and Protease and Phosphatase Inhibitor Cocktail (Cat#89901, Thermofisher Scientific). Lysed samples were agitated and centrifuged at 4°C. Supernatant was aliquoted and stored at-80°C. Protein concentration was obtained using protocol from 5.2 Microplate Assay Protocol of Detergent Compatible (DC) Protein Assay Instruction Manual (BioRad). FlexStation 3 (Molecular Devices) was used to measure protein concentration at 750nm.

Prior to loading samples, protein was denatured at 95°C for 5 minutes with 6x Laemmli SDS buffer (Cat#J61337-AD, Thermofisher Scientific). Samples were loaded into a 10% Mini-PROTEAN TGX Gel (Cat#4561034, BioRad) and run at 150V until separated. Protein was transferred to a PVDF membrane (Cat#1620177, BioRad) for 45 minutes at 100V. The PVDF membrane was blocked using Intercept Blocking Buffer (Cat#927-60001, LI-COR) for 1 hour at room temperature and incubated with primary antibodies overnight in Intercept Antibody Diluent (Cat#927-65001, LI-COR) overnight at 4°C. Primary antibodies are as follows: Rb LRP2 (Cat#AB76969, Abcam, 1:400), and Ms β-actin (Cat#3700, Cell Signaling Technology, 1:1500). After overnight incubation in primary, PVDF membrane was washed 3x with TBS-T before incubation in the secondary antibody. Secondary antibodies are as follows: IRDye 680RD Goat anti-Mouse IgG (Cat#926-68070, LI-COR, 1:10,000), IRDye 800CW Donkey anti-Rabbit (Cat#926-32213, LI-COR, 1:10,000).

Secondaries are diluted in Antibody Diluent for one hour at room temperature, after, the membrane is washed 3x with TBS-T, and a final wash in TBS. Membrane was imaged on the Odyssey CLx (LI-COR) using ImageStudio Ver 5.2 (LI-COR). ImageJ and GraphPad Prism version 10.1.2 were used for data analysis.

### TUNEL

Paraffin-embedded kidney sections were deparaffinized and washed twice in PBS. Antigen retrieval was performed by incubating sections with Proteinase K for 30 minutes at room temperature (RT), followed by two PBS washes. The TUNEL reaction was carried out using the TUNEL Assay Kit, fluorescence 594 nm (Cat#48513, Cell Signaling Technology), according to the manufacturer’s instructions. After a final wash with PBS-TB, sections were incubated overnight at 4 °C with goat anti-rat KIM-1 antibody (Cat#AF3689, R&D Systems). The following day, sections were washed in PBS-T and incubated with donkey anti-goat Alexa Fluor 750 secondary antibody, then mounted with Fluoromount-G™ Mounting Medium (Cat#5018788, Fisher Scientific). Imaging was performed using a Nikon Ti2 inverted widefield microscope with a SpectraX light engine at 20x magnification. Image analysis was conducted using NIS-Elements AR 6.20.02 software. For quantification, the kidney cortex was manually delineated on whole-section images, and a GA3 analysis recipe was created to measure: (1) the number of TUNEL-positive puncta, (2) total TUNEL-positive area, (3) KIM1-positive area within the cortex, and (4) total cortical area.

### Quantitative Proteomic analysis of LRP2

Four animals (two male, two female) were analyzed at P4, P10, and adulthood with 5 days of toxic gent or saline exposure (n = 24 total). Kidneys were collected 3 hours after 4^th^ dose. Kidneys were snap frozen in ethanol and dry ice. Membrane proteins were extracted from rat kidney cortex (∼30–60 mg) using the Thermo Scientific™ Mem-PER™ Plus Membrane Protein Extraction Kit, followed by data acquisition using LC-MS-based quantitative proteomics. LRP2 abundance was quantified using Total Proteomics Approach (TPA) and the data were normalized to total membrane protein abundance as previously described.(62, 63) Additional details can be found in the **Supplemental Methods**.

### Lectin labeling

Lycopersicon Esculentum (Tomato) Lectin (LEL, TL), DyLight® 649 (DL-1178-1, 1 ug/g body weight) was administered to rats under Triple Sedative Solution NON-Survival Procedures (67 mg/mL Ketamine + 3.3 mg/mL Xylazine + 1.7 mg/mL Acepromazine) and isoflurane anesthesia via cardiac injection. After 5 minutes of circulation, animals underwent terminal perfusion as previously described(17) using 30ml PBS and 30ml of 4% paraformaldehyde (PFA). This protocol was modified from Kinoshita et al.(64)

### Clearing

Kidneys were fixed in PFA and subsequently washed twice with phosphate-buffered saline (PBS). Following fixation, tissues were dehydrated in a graded series of tetrahydrofuran (THF)/PBS solutions at 50%, 75%, 90%, and 100% for 2 hours each, followed by an additional overnight incubation in 100% THF. Delipidation was performed using 100% dichloromethane (DCM) for an initial 2-hour incubation and a subsequent 24-hour incubation. The kidneys were then rehydrated through a reverse THF/water gradient consisting of 100%, 90%, 75%, and 50% THF, and finally transferred to 100% water overnight. Rehydrated kidneys underwent heme extraction in a solution containing 25% Quadrol and 5% CHAPS for 4 days at 37 °C on a rocking platform. Following heme extraction, the tissues were dehydrated again through a graded methanol series (25%, 50%, 75%, 90%, and 100%), with each step lasting 2.5 hours. This was followed by two additional changes in 100% methanol over 48 hours. Finally, the kidneys were incubated in benzyl alcohol-benzyl benzoate (BA:BB) mixture (1:2) overnight and transferred to fresh BA:BB solution the following day and incubated on a rocker for at least 48 hours before imaging.

### Light sheet imaging

All lectin perfused kidneys were imaged on the Bruker LCS Spim light sheet and processed using LuxBundle. Both calibration and imaging had a camera exposure time of 50ms and 1ms. Calibration was done prior to imaging each kidney to account of differences in the refractive index of BA:BB on illumination channel 488nm with an exposure of 60%, and detection filter wheel 498. During calibration, scanner settings were offset at-0.1 (left scanner) and +0.1 (right scanner), beams (OL and OR) were adjusted across 3 points on the Y axis. After calibration, the kidney was placed in the cuvette and submerged in BA:BB. Illumination was switched to 685nm with an exposure of 80%, uniform illumination amplitude was enabled at 45%, and switched to detection filter wheel LP 700. OZ settings were adjusted to increase focus and added across several planes of the kidney containing high number of glomeruli. A stack was created to include the entire kidney with a step size of 2700, stack was added as an event to the experiment tab and imaged at 2.2x. Once complete, completed image was added to the processing tab and stitched to obtain the final Imaris file.

### Nephron Quantification from light sheet images

The light sheet images were used to quantify volume and number of nephrons using analysis software Imaris 11.0.0 and NIS Elements AR 6.20.02. For nephron counts, Imaris files obtained from the light sheet were converted to.nds format and were clarified and denoised. The files were then converted back to.ims using Imaris File Converter 11.0.1. In Imaris a workflow was created to identify spots (lectin labelled nephrons) and used machine learning to segment nephrons from other artifacts. The workflow was ran multiple times to be trained and used the version that best suited all samples. The workflow was then run on all samples and nephron counts were exported.

For nephron volumes, a workflow was created in which the whole kidney surface was rendered using absolute intensity thresholding. Then a second “Surface creation” module is added to the workflow to render individual nephrons using the machine learning segmentation method. The training data is saved within the workflow and multiple runs were done to train the segmentation process to accurately determine nephron volumes across different samples. The final workflow was then used to determine nephron volumes and the data was exported. 11 samples were insufficiently cleared for light sheet imaging: F 9E, R 3E, SS 4C, SS 8C, SS 10C, TS 4C, TS 9E, TS 12C, VC 8C, WS 9E, and WS 11E. However, all samples were included in glomerular density automated quantification (see below).

### Trichrome

Masson’s Trichrome staining was performed on 5µm paraffin embedded kidney sections by the pathology core at Cincinnati Children’s Hospital. Images were captured on the Nikon NiE Upright Microscope at 10x magnification and processed using NIS-Elements software, version AR 6.20.02. Semiquantitative scoring was performed by a clinical pathologist blinded to experimental decisions. A scoring system was developed where staining was categorized by focality and then gave a severity score on a 0-3 scale, with 0 having no fibrosis and 3 being the most severe.

### Glomerular Density automated quantification

Whole-slide trichrome images were converted from Nikon ND2 format into pyramidal OME-TIFF files for downstream analysis. 4-fold reductions (2.36 μm/pixel) were extracted and used for downstream processing. Renal cortex regions were manually annotated in QuPath (v0.5.1) to exclude noncortical tissue from analysis. Glomerular segmentation was performed using a U-Net convolutional neural network trained to identify both normal and abnormal glomeruli, including Bowman’s capsule.(65) Segmentation performance was validated against annotations from two independent observers with F1 scores of 95.69% and 92.51%, respectively.

Segmented glomerular masks and cortex regions of interest (ROI) annotations were subsequently analyzed in Fiji/ImageJ (v1.54t). For each image, cortical area, glomerular count, and glomerular density (glomeruli/mm^2^ cortex) were calculated. Full methods can be found in **Supplemental Methods**.

### RT-qPCR For P4 Rat Sex Determination

RT-qPCR for P4 rat sex determination was completed as previously described.(66) Presence of *SRY* gene primer(67) (Sigma) was used to determine sex. *Gapdh*(68) and *Ppia* were used as housekeeping genes, no normalization was required. Primer sequences found in **Supplemental Table 5**. RT-qPCR results in **Supplemental Table 6**. Two adult male and female rats were used as positive and negative controls to confirm methodology.

## Statistical Analyses

Using data from a previous rabbit study(17), sample size calculations were performed for a two-sample t-test assuming a 30% difference in mean nephron number between the saline and gent groups, a coefficient of variation (SD/mean) between animal of 10%, a two-sided α of 0.05, and 80% power. Under these assumptions, the estimated sample size was approximately 3 animals per group. We therefore targeted 3–4 animals for each combination of prenatal treatment, postnatal treatment, and period. Statistical analyses were performed using GraphPad Prism v9 and SAS 9.4. All animals were used regardless of sex. Descriptive data were calculated for glomerular counts, glomerular volume, BUN, and sCR. The normality of the data was assessed using Shapiro-Wilk test before using parametric vs nonparametric comparisons. Differences in BUN, PCR, and nephron number were assessed with one-way ANOVA with multiple comparisons (Tukey’s) if normally distributed, or Kruskal-Wallis with Dunn’s multiple comparisons if not normally distributed. RT-qPCR data were found to be normally distributed and comparisons were made using unpaired t-tests on the delta Ct. Kidney weights and kidney-to-body weights did not pass normality testing and were analyzed with Kruskal-Wallis testing. When comparing nephron numbers, controls and beta without gentamicin exposures were combined.

### Random-intercept linear mixed-effects model

Random-intercept linear mixed-effects models were fitted, with litter included as a random effect. Glomerular count and glomerular density were respectively evaluated as response variables. For each outcome, fixed effects included prenatal treatment, postnatal period, and their interaction. Prenatal treatment was categorized as beta or no prenatal treatment. Postnatal treatment and treatment period were combined into a single three-level variable, Postnatal period, with levels of Gent P0–4, Gent P6–10, and Saline P0–4 or P6–10. Six prespecified pairwise comparisons were performed for each outcome. P-values for the pairwise comparisons were adjusted for multiple comparisons using the Bonferroni method. Estimated differences and adjusted P-values are presented in Tables 3.

## Study approval

This study was reviewed and approved by the Cincinnati Children’s Hospital Medical Center (CCHMC) Institutional Animal Care and Use Committee (IACUC) under protocols 2022-0042 and 2025-0036.

## Data availability

All data will be made available upon request. Data processing is listed in detail in methods. Data presented in figures is available in supplemental tables.

## Author contributions

MPS conceptualized the study. CN, BB, SY performed the majority of the experiments. CN performed the light sheet imaging. SY performed the light sheet quantification. KS helped with breeding an experimental set up of the animal colony. SI helped with the experiments and acquired and analyzed data. KV performed the semiquantitative analyses of trichrome staining. JB performed the glomerular density analyses on trichrome staining. AV and JER assisted with western blot experiments. DS and BP performed the proteomic analyses. CN and MPS wrote the first draft of the manuscript, and all authors edited the final version of the manuscript.

## Acknowledgments

This research was made possible, in part, by using the Bio-Imaging and Analysis Facility (RRID:SCR_022628), Integrated Pathology Research Facility (RRID:SCR_022637), and the Comparative Medicine Facility. We specifically want to thank Drs. Kentaro Iwasawa and Matthew Kofron for their guidance in nephron labeling and quantification using light sheet microscopy.

