## Supplemental file for "Megalin (LRP2), prenatal betamethasone, and injury susceptibility in the developing kidney"

**Supplemental file Table of Contents:**

1) Supplemental Methods: 1-6

2) Supplemental Figures: 7-12

3) Supplemental Tables: 13-24

4) Supplemental Movie: 25

**Supplemental Methods:**

**Image preprocessing for glomerular density**

Whole-slide trichrome images were preprocessed from Nikon ND2 format into pyramidal OME-TIFF files for downstream analysis. ND2 files were batch-submitted on the HPC cluster, with one Slurm worker job launched per .nd2 file. Each ND2 image was read using nd2, converted to a NumPy array, squeezed to remove singleton dimensions, and saved as a BigTIFF pyramidal OME-TIFF using tifffile. Image pyramids were generated by repeated 2× spatial decimation, producing the original full-resolution image plus four progressively downsampled levels. The resulting files were saved with the suffix _pyramidal.ome.tiff.

To generate reduced-resolution TIFF images for model input and downstream processing, the pyramidal OME-TIFF files were then reopened using tifffile, and the stored pyramid levels were extracted. Level 2 was exported as the 4-fold reduced image at 2.36 μm/pixel aspect ratio. These outputs were saved into separate 4fold_reduced, while preserving the original image stem in the output filename.

4-fold reduced images were converted to flattened RGB Color images for U-Net Segmentation processing.

Cortex outlines were manually performed in QuPath (version 0.5.1) for each flattened 4-fold reduced trichrome image to identify the renal cortex region and exclude non-cortical tissue from downstream analysis. These annotated cortex regions were then exported and used for subsequent preprocessing and image analysis workflows.

**U-Net Model Segmentation**

The U-Net convolutional neural network architecture was used for all image segmentation tasks. The model was trained on 500x500 pixel images from the 4-fold down-sampled trichrome images containing glomeruli that were manually annotated in QuPath. We trained a model for glomerular segmentation using the 4-fold down-sampled images to include each individual glomerulus with its accompanying Bowman's Capsule. The model was trained to identify both normal and abnormal appearing glomeruli.

The results of this analysis were validated by comparing the U-Net-derived glomerular identification to those obtained by two independent human observers using a develop quality control code that identified two

650x650 pixel images from each trichrome image and requested the user to input the number of properly detected glomeruli (true positives), non-glomeruli detected (false positives), and non-detected glomeruli (false negatives). Recall, Precision, and F1 scores, defined below, were calculated based on true positives (TP), false positives (FP), and false negatives (FN) identified.

$$Recall = \frac{TP}{TP + FN}$$

$$Precision = \frac{TP}{TP + FP}$$

$$F1 = \frac{2 \times Precision \times Recall}{Precision + Recall}$$

Recall, precision, and F1 scores were evaluated using two 650x650 pixel regions sampled from each of 38 trichrome images, resulting in a total of 76 image sections assessed by two independent validators. Validator 1 achieved recall, precision, and F1 scores of 95.38%, 97.04%, and 95.69%, respectively, while Validator 2 achieved scores of 91.38%, 95.75%, and 92.51%, respectively.

**Post Image Analysis**

Segmented glomeruli masks and corresponding cortex ROIs were analyzed in ImageJ/Fiji (Version 1.54t 16 May 2026) using a custom macro. For each image, the segmented glomeruli image was opened, converted to a binary mask, and paired with its manually generated cortex outline. The cortex ROI was added to the ROI Manager and measured to obtain total cortical area. Glomeruli were then counted within the cortex ROI using particle analysis with minimum size threshold of 1000 $\mu m^2$ to exclude small artifacts. For each kidney image, the macro recorded the image ID, glomeruli count, cortex area, and glomerular density, calculated as glomeruli count divided by cortex area. Results were exported as a timestamped CSV file for downstream statistical analysis.

**Tissue homogenization and total membrane protein isolation:**

Total membrane proteins were extracted from rat kidney tissue homogenates using the Mem-PER™ Plus kit (Product Code: 89842). Prior to homogenization, the required volume of permeabilization buffer supplemented with 1% protease inhibitor was prepared. Each pre-weighed kidney tissue sample was washed with approximately 2 mL of cell wash solution, briefly vortexed, and the wash was discarded. Juvenile and adult rat kidney tissues were homogenized in ice-cold permeabilization buffer containing 1% protease inhibitor, maintaining a tissue (mg)-to-buffer (mL) ratio of 100:1. Homogenization was performed using a bead homogenizer (VWR, Radnor, PA) for four cycles of 15 seconds each at a speed setting of 2 (arbitrary units), with samples placed on ice for approximately 30 seconds between cycles. Following homogenization, all samples were incubated at 4 °C for 10 minutes with gentle shaking at 300 rpm. The homogenates were then centrifuged at 1,500 × g and 4 °C for 10 minutes to remove the nuclear fraction. The resulting supernatant was collected and further incubated at 4 °C for 30 minutes with gentle shaking, followed by centrifugation at 16,000 × g and 4 °C for 15 minutes. The supernatant containing cytosolic proteins was transferred to new pre-labeled tubes. The remaining pellets were resuspended in membrane solubilization buffer (MEM-PER solubilization buffer containing 4% SDS, diluted 1:1, and supplemented with 1% protease inhibitor). The samples were then incubated at 15 °C for 60 minutes with gentle shaking at 300 rpm to facilitate membrane protein solubilization. The solubilized total membrane protein fractions were stored at −80 °C until further use. Protein concentrations were determined using a BCA assay kit according to the manufacturer's instructions.

**Protein digestion**

The processed kidney samples were diluted to a protein concentration of 1 mg/mL using a membrane solubilization buffer, prior to digestion. The protein sample was then denatured, alkylated, desalted, and digested as per in-house optimized digestion protocol. Briefly, 80 µg of protein in 80 µL was denatured and reduced by heating at 95°C for 10 minutes after mixing with 10 µL of BSA (0.2 mg/mL), 20 µL of DTT (250 mM), and 30 µL of ammonium bicarbonate (100 mM). The sample was cooled to room temperature for 10 minutes, followed by alkylation with 10 µL of freshly prepared IAA (500 mM) in 100 mM ammonium bicarbonate. The sample was incubated at room temperature in the dark for 30 minutes. Alkylated proteins were precipitated by adding 1 mL of ice-cold acetone and incubating for 2 hours at −80°C. The precipitated protein sample was centrifuged at 16,000 × g for 15 minutes at 4°C, and the supernatant was carefully discarded. The protein pellet was washed with 500 µL of ice-cold methanol and centrifuged again at 16,000 × g for 15 minutes at 4°C. After removal of the supernatant, the pellet was dried for 30 minutes. The dried protein pellet was dissolved in 60 µL of 50 mM ammonium bicarbonate solution, followed by the addition of 20 µL of trypsin solution (0.16 µg/µL, freshly prepared in 50 mM ammonium bicarbonate buffer), and incubated at 37°C for 16 hours with shaking at 350 rpm. The protein-to-trypsin ratio was 25:1 (w/w). Digestion was quenched by adding 5 µL of 5% formic acid solution, and the samples were centrifuged at 16,000 × g for 15 minutes prior to LC-MS analysis.

**Mass spectrometry data acquisition**

Global proteomics data were acquired using an EASY-Spray 1200 series nanoLC system coupled to a Q Exactive HF mass spectrometer (Thermo Fisher Scientific, Waltham, MA), equipped with a Thermo Scientific Acclaim PepMap™ 100 pre-column (75 µm × 2 cm, Part No. 164535) and a Thermo Scientific PepMap™ RSLC C18 analytical column (75 µm × 25 cm, Part No. ES902). The mobile phases consisted of 0.1% formic acid in water (A) and 0.1% formic acid in 80% acetonitrile (B). Peptides were eluted using the following gradient (%B): 0–10% (0–2 min), 10–45% (2–27 min), 45–100% (27–28 min), and 100% (28–35 min). The flow rate and column temperature were maintained at 300 nL/min and 40 °C, respectively. Proteomics data were acquired with a 1 µL injection volume (corresponding to 1 µg of protein digest). Full MS survey scans (m/z 348–1100) were acquired with an automatic gain control (AGC) target of $3 \times 10^6$, a resolution of 60,000, and a maximum ion injection time of 55 ms. Data were collected in data-independent acquisition (DIA) mode at a resolution of 30,000, with a maximum ion injection time of 55 ms and a normalized collision energy of 30 eV in positive ionization mode. DIA data were acquired using variable isolation windows as follows: m/z 350–400 (25 m/z window), m/z 400–870 (20 m/z window), and m/z 870–1110 (40 m/z window). The spray voltage, capillary (ion transfer tube) temperature, and S-lens RF level were set to 1.7 kV, 300 °C, and 50 (arbitrary units), respectively.

**Quantitative proteomics analysis of LRP2**

After MS acquisition, DIA raw data were processed using Data-Independent Acquisition by Neural Networks (DIA-NN) software (version 1.8.1). FASTA files for *Rattus norvegicus* (rat) were downloaded from the UniProt Knowledgebase (UniProtKB) for spectral library generation. Peptide identification was performed using *in silico* digestion with trypsin specificity (cleavage at K* and R*), allowing up to two missed cleavages. Peptides with lengths between 7 and 30 amino acids were considered. Precursor ion m/z values were restricted to a range of 300–1800, with charge states from 1 to 4, and fragment ions were filtered within an m/z range of 200–1800.

Carbamidomethylation of cysteine was set as a fixed modification, while oxidation of methionine was included as a variable modification. Quantification was performed using the fixed-width center of each elution peak. Highly heuristic protein grouping was applied to reduce the number of inferred protein groups, and RT-dependent cross-run normalization with match-between-runs (MBR) was enabled. The output was filtered at a 1% false discovery rate (FDR). Protein abundance of LRP2 was determined using the total protein approach (TPA), followed by normalization to a marker protein.

**Supplemental Figure legends**

**Supplemental Figure 1: Postnatal rat nephrogenesis.** Representative immunofluorescence imaging demonstrating developmental expression patterns of rat nephrogenesis using SIX2, SOX9, CDH6, LRP2, ZO-1, and KRT8 across several postnatal timepoints.

**Supplemental Figure 2: Experimental design for exposures**

**Supplemental Figure 3: KIM1 Injury Staining at P4 and P10. A-B)** Increased KIM1 staining seen at cortico-medullary border of P10 kidneys compared to P4. **C)** Increased KIM1 staining in beta + gent P6-10 compared to gent P6-10 only seen in B.

**Supplemental Figure 4: Prenatal betamethasone and early (P0-4) gentamicin exposure: No difference in long-term outcomes. A)** There was a trend of increased BUN in prenatal beta (23mg/dL) and beta + gent P0-4 (23.17mg/dL), relative to control (19.4mg/dL) or gent P0-4 (20.8mg/dL). However, no significant difference was identified after multiple comparisons. B-C) No significant difference in PCr seen in between P0-4 exposure groups ($p = 0.538$) or nephron number ($p = 0.719$). D-E) Histological trichrome analyses from prenatal betamethasone treated kidneys (D, $D^1$) and the beta + gent P0-4 (E, $E^1$) treatment group in adulthood.

**Supplemental Figure 5: LRP2 Western blot for early and late controls with and without prenatal betamethasone. A)** Western blot analysis and quantification of LRP2 protein expression in P10 beta vs control identified significantly increased in LRP2 at P10 after prenatal beta exposure with **B)** representative western blot images of LRP2 and β-Actin. **C)** Western blot analysis and quantification of LRP2 protein expression in P4 beta vs control showed no difference in LRP2 at P4 ($p = 0.886$) with **D)** representative western blot images of LRP2 and β-Actin. Both plots depict median with IQR.

Supplemental Figure 1

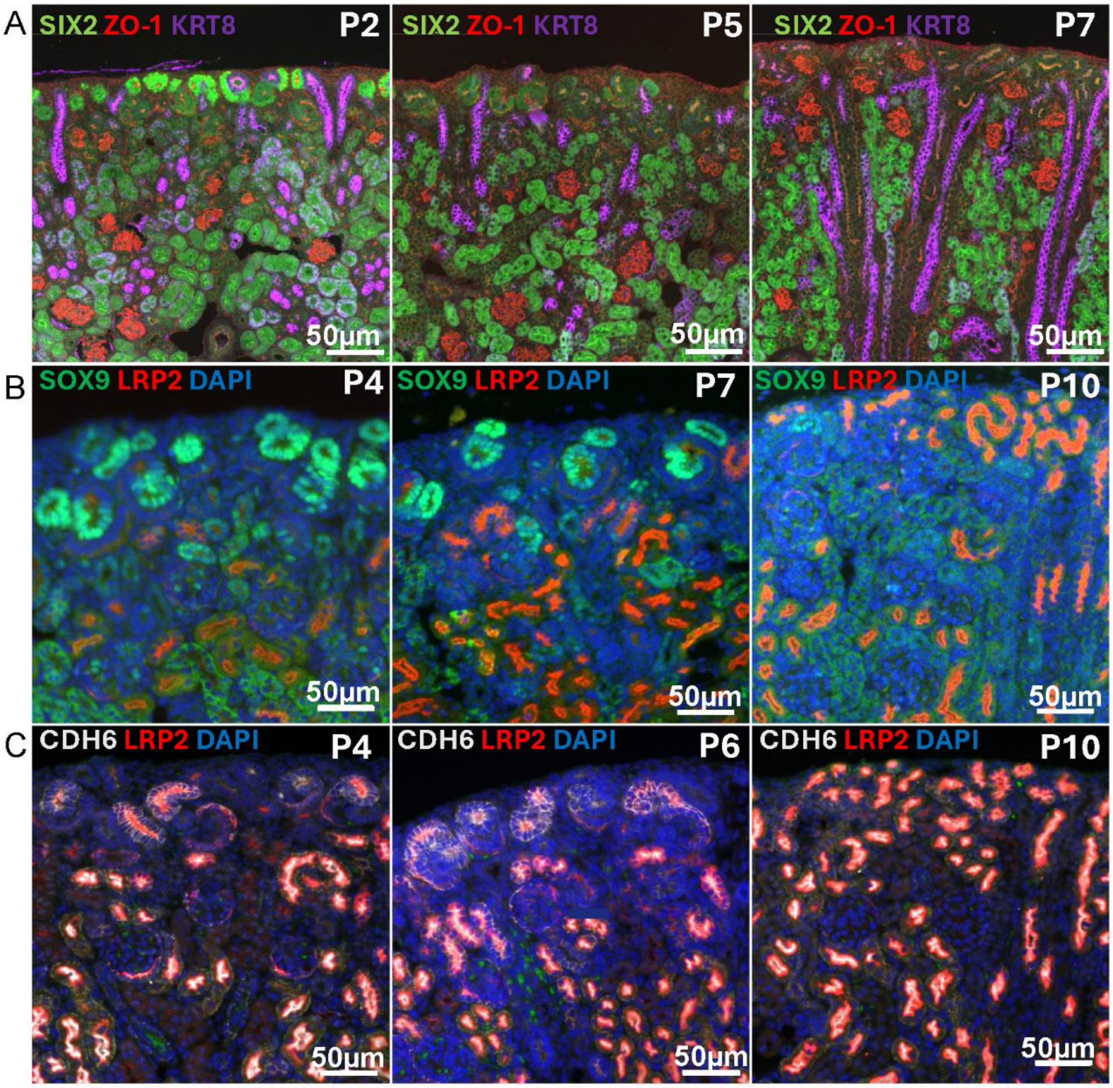

Supplemental Figure 2

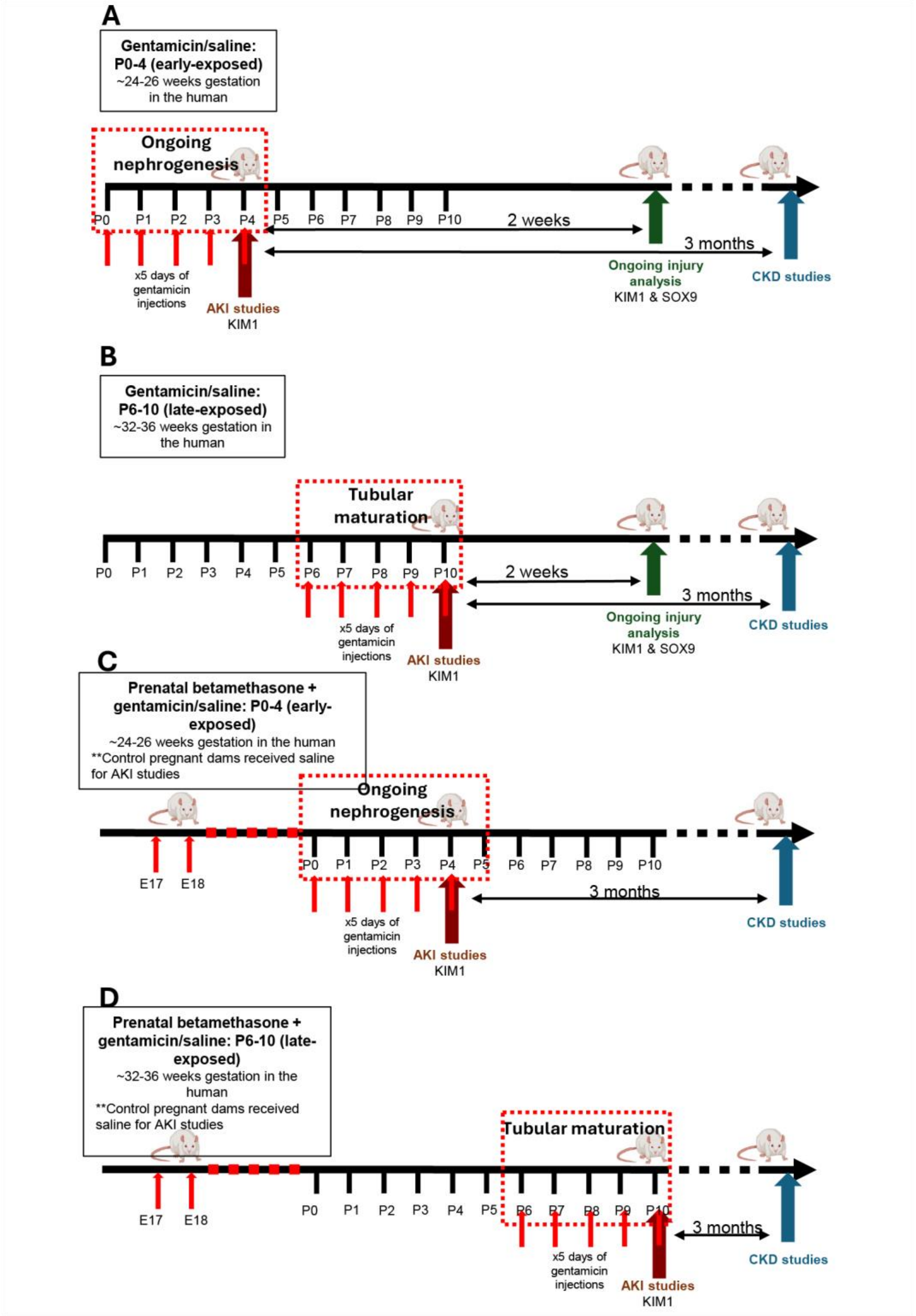

Supplemental Figure 3

A) Gent P0-P4 KIM1 LRP2 DAPI

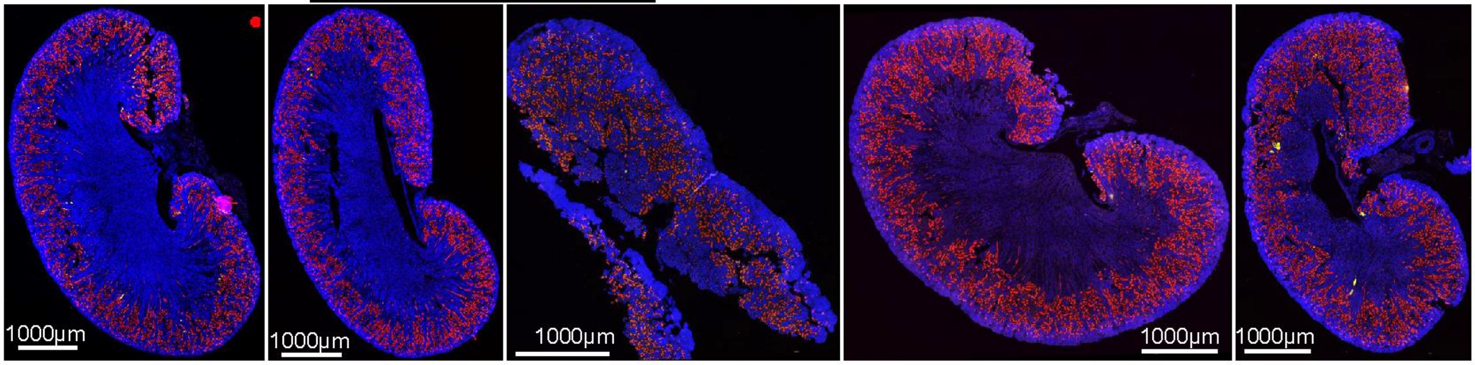


B) Gent P6-P10 KIM1 LRP2 DAPI

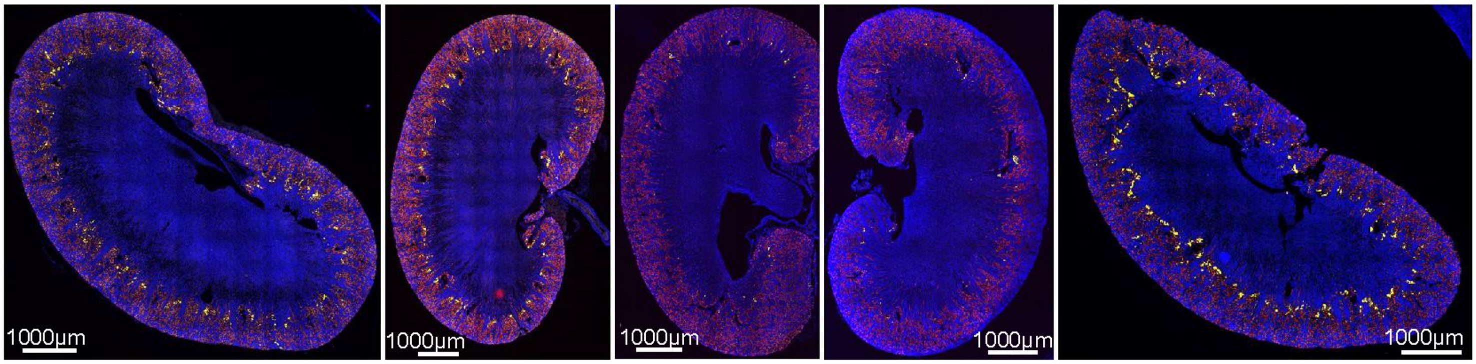


C) Beta + Gent P6-P10 KIM1 LRP2 DAPI

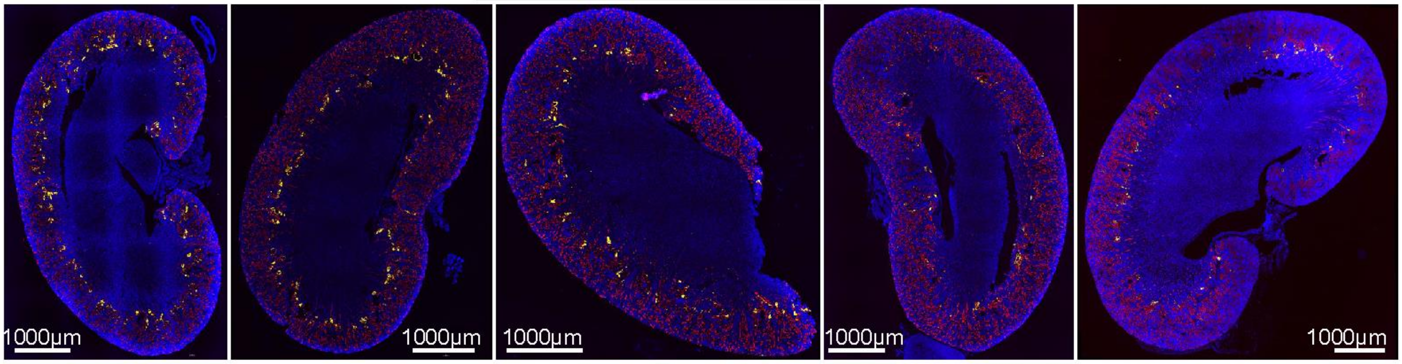

Supplemental Figure 4

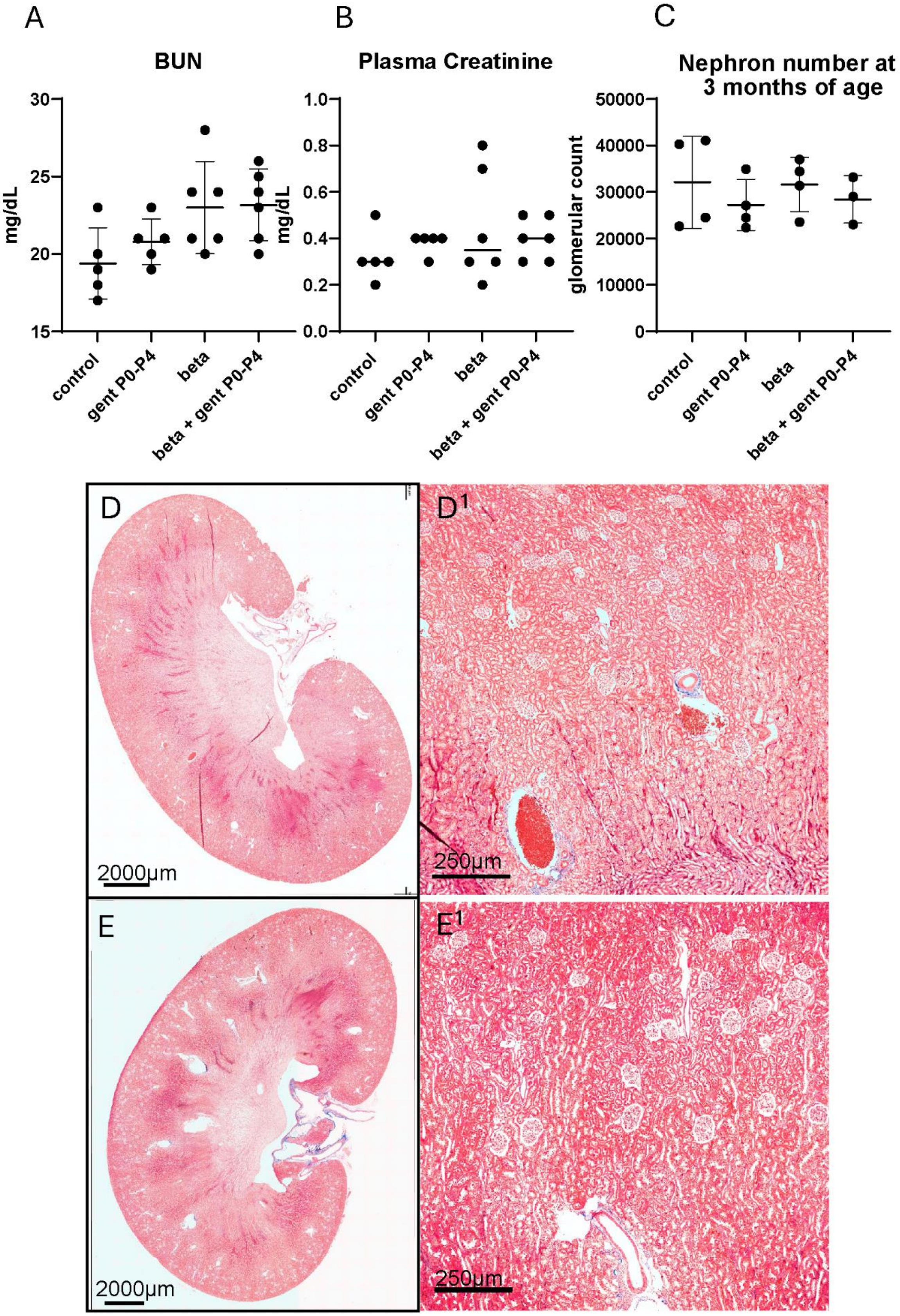

Supplemental Figure 5

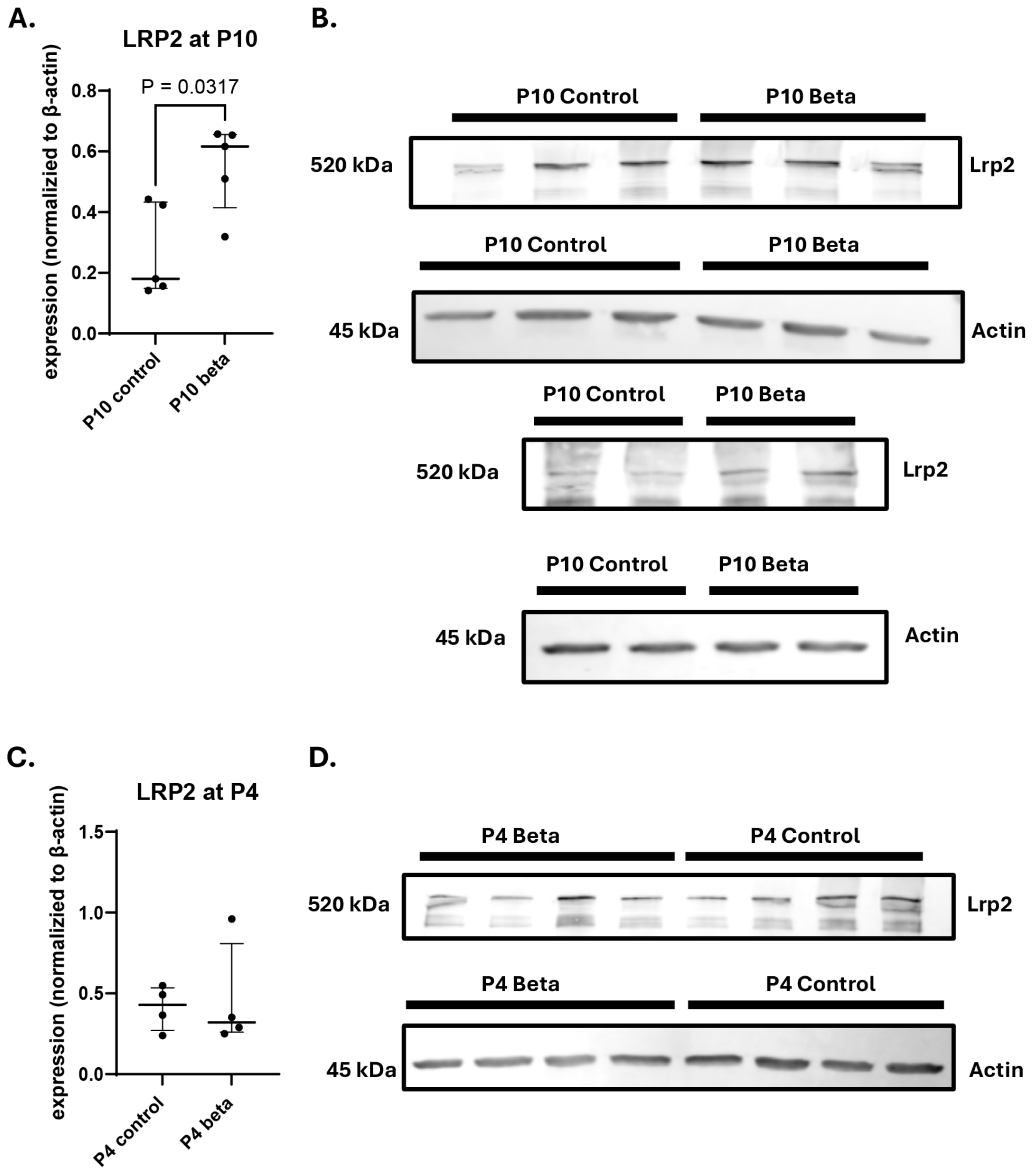

**Supplemental Table 1: Semiquantitative scoring system of fibrosis.** Semiquantitative fibrosis scoring for all adulthood experimental groups, including regional distribution of fibrosis scarring identified during analysis by clinical pathologist.

**Supplemental Table 2: Glomerular density.**

**Supplemental Table 3: Glomerular volume for all adulthood samples.** Glomerular volume measurements for each adult experiment group, including sample size, average glomerular volume, variability, and distribution of analyzed glomeruli.

**Supplemental Table 4: All samples used for this study with exposures and treatments.** Sample information for all animals included in the study, including experimental group assignment, biological sex, litter origin/name, collection timepoint, and treatments administered.

**Supplemental Table 5: Primer sequences used for qPCR and sex-determination qPCR.** Forward and reverse primer sequences used for RT-qPCR experiments, including target genes, and housekeeping genes.

**Supplemental Table 1**

| Treatment | Sample | IDs | Fibrosis score | Location | Focality |
|---|---|---|---|---|---|
| P0-4 control | R11025 | 1E | 1 | Inner cortex | Focal |
| | | 3E | 1 | Inner cortex | Focal |
| | | 5E | 1 | Inner cortex | Focal |
| | | 7E | 1 | Inner cortex | Focal |
| | | 9E | 1 | Inner cortex | Focal |
| | | 2C | 1 | Inner cortex | Focal |
| | | 4C | 1 | Inner cortex | Focal |
| | | 6C | 1 | Inner cortex | Focal |
| | | 8C | 0 | None | None |
| | | 10C | 1 | Inner cortex | Focal |
| P0-4 steroid | TS21825 | 1E | 1 | Inner cortex | Focal |
| | | 7E | 2 | Inner to mid cortex | Focal |
| | | 9E | 1 | Inner cortex | Focal |
| | | 13E | 1 | Inner cortex | Focal |
| | | 2C | 0 | None | None |
| | | 4C | 1 | Inner cortex | Focal |
| | | 10C | 1 | Inner cortex | Focal |
| | | 12C | 1 | Inner cortex | Focal |
| P6-10 steroid | SS20525 | 3E | 3 | Inner to outer cortex | Diffuse |
| | | 7E | 3 | Inner to outer cortex | Diffuse |
| | | 9E | 2 | Inner to outer cortex | Diffuse |
| | | 11E | 3 | Inner to outer cortex | Diffuse |
| | | 4C | 1 | Inner cortex | Focal |
| | | 6C | 1 | Inner cortex | Focal |
| | | 8C | 1 | Inner cortex | Focal |
| | | 10C | 1 | Inner cortex | Focal |
| P6-10 control | F91224 | 3E | 1 | Inner cortex | Focal |
| | | 5E | 1 | Inner cortex | Focal |
| | | 9E | 2 | Inner to mid cortex | Diffuse |
| | | 15E | 1 | Outer cortex | Focal |
| | | 6C | 1 | Inner cortex | Focal |
| | | 10C | 0 | None | None |
| | | 14C | 0 | None | None |
| | | 16C | 0 | None | None |

Fibrosis- 0-3 scale (None, mild, <10%, moderate- 10-50%, severe >50% cortex involved)

### Supplemental Table 2

| Prenatal treatment | Postnatal treatment | Treatment period | Sample ID | Image ID | Glomeruli count | Cortex area (mm^2) | Glomerular density (1/mm^2) |
|---|---|---|---|---|---|---|---|
| None | Saline | P6-10 | F91224-10C | F91224-LF-10C_pyramidal_4x_SEGMENTATION.tif | 578 | 135.1812375 | 4.275741297 |
| None | Saline | P6-10 | F91224-14C | F91224-LF-14C_pyramidal_4x_SEGMENTATION.tif | 432 | 100.6799326 | 4.290825279 |
| None | Gent | P6-10 | F91224-15E | F91224-LF-15E_pyramidal_4x_SEGMENTATION.tif | 440 | 75.34919132 | 5.83947873 |
| None | Gent | P6-10 | F91224-3E | F91224-LF-3E_pyramidal_4x_SEGMENTATION.tif | 462 | 74.90086637 | 6.168152952 |
| None | Gent | P6-10 | F91224-5E | F91224-LF-5E_pyramidal_4x_SEGMENTATION.tif | 405 | 84.11696146 | 4.814724557 |
| None | Saline | P6-10 | F91224-6C | F91224-LF-6C_pyramidal_4x_SEGMENTATION.tif | 472 | 70.09951471 | 6.733284844 |
| None | Gent | P6-10 | F91224-9E | F91224-LF-9E_pyramidal_4x_SEGMENTATION.tif | 358 | 104.152517 | 3.437266909 |
| Saline | Saline | P6-10 | VC31925-8C | VC31925_8C_pyramidal_4x_SEGMENTATION.tif | 409 | 122.984894 | 3.325611679 |
| Beta | Saline | P6-10 | SS20525-10C | SS20525_10C-M_trichrome_pyramidal_4x_SEGMENTATION.tif | 370 | 139.1946412 | 2.658148309 |
| Beta | Gent | P6-10 | SS20525-11E | SS20525_11E-M_trichrome_pyramidal_4x_SEGMENTATION.tif | 277 | 110.6171405 | 2.504132711 |
| Beta | Gent | P6-10 | SS20525-3E | SS20525_3E-F_trichrome_pyramidal_4x_SEGMENTATION.tif | 269 | 73.24453416 | 3.672628997 |
| Beta | Saline | P6-10 | SS20525-4C | SS20525_4C-F_trichrome_pyramidal_4x_SEGMENTATION.tif | 396 | 75.33218733 | 5.256717136 |
| Beta | Saline | P6-10 | SS20525-6C | SS20525_6C-M_trichrome_pyramidal_4x_SEGMENTATION.tif | 424 | 99.66731251 | 4.254153035 |
| Beta | Gent | P6-10 | SS20525-7E | SS20525_7E-M_trichrome_pyramidal_4x_SEGMENTATION.tif | 236 | 94.04971927 | 2.509311052 |
| Beta | Saline | P6-10 | SS20525-8C | SS20525_8C-M_trichrome_pyramidal_4x_SEGMENTATION.tif | 391 | 106.305702 | 3.678071754 |
| Beta | Gent | P6-10 | SS20525-9E | SS20525_9E-M_trichrome_pyramidal_4x_SEGMENTATION.tif | 312 | 125.3576662 | 2.488878499 |
| None | Gent | P0-4 | R11025-1E | R11025-1E_pyramidal_4x_SEGMENTATION.tif | 583 | 106.20091 | 5.489595145 |
| None | Saline | P0-4 | R11025-2C | R11025-2C_pyramidal_4x_SEGMENTATION.tif | 423 | 86.17718435 | 4.908491768 |
| None | Gent | P0-4 | R11025-3E | R11025-3E_pyramidal_4x_SEGMENTATION.tif | 517 | 87.99855494 | 5.875096476 |
| None | Saline | P0-4 | R11025-4C | R11025-4C_pyramidal_4x_SEGMENTATION.tif | 529 | 96.59101078 | 5.476700116 |
| None | Gent | P0-4 | R11025-5E | R11025-5E_pyramidal_4x_SEGMENTATION.tif | 487 | 91.88528916 | 5.300086711 |
| None | Saline | P0-4 | R11025-6C | R11025-6C_pyramidal_4x_SEGMENTATION.tif | 410 | 102.7755058 | 3.989277377 |
| None | Gent | P0-4 | R11025-7E | R11025-7E_pyramidal_4x_SEGMENTATION.tif | 430 | 60.02307237 | 7.16391186 |
| None | Saline | P0-4 | R11025-8C | R11025-8C_pyramidal_4x_SEGMENTATION.tif | 449 | 71.97982838 | 6.237858718 |
| None | Gent | P0-4 | R11025-9E | R11025-9E_pyramidal_4x_SEGMENTATION.tif | 404 | 50.71693912 | 7.965780408 |
| Beta | Saline | P0-4 | TS21825-10C | TS21825_trichrome_10C-M-1_pyramidal_4x_SEGMENTATION.tif | 442 | 140.3086002 | 3.150198915 |
| Beta | Saline | P0-4 | TS21825-12C | TS21825_trichrome_12C-F-1_pyramidal_4x_SEGMENTATION.tif | 385 | 70.61579435 | 5.452038082 |
| Beta | Gent | P0-4 | TS21825-13E | TS21825_trichrome_13E-M-1_pyramidal_4x_SEGMENTATION.tif | 459 | 102.103411 | 4.495442371 |
| Beta | Gent | P0-4 | TS21825-1E | TS21825_trichrome_1E-F-1-redo_pyramidal_4x_SEGMENTATION.tif | 408 | 83.74603724 | 4.871872311 |
| Beta | Saline | P0-4 | TS21825-2C | TS21825_trichrome_2C-F-1_pyramidal_4x_SEGMENTATION.tif | 448 | 76.63065046 | 5.846224681 |
| Beta | Saline | P0-4 | TS21825-4C | TS21825_trichrome_4C-M-1_pyramidal_4x_SEGMENTATION.tif | 560 | 114.2110085 | 4.903205106 |
| Beta | Gent | P0-4 | TS21825-7E | TS21825_trichrome_7E-M-1-redo_pyramidal_4x_SEGMENTATION.tif | 427 | 121.6826714 | 3.509127429 |
| Beta | Gent | P0-4 | TS21825-9E | TS21825_trichrome_9E-M-1_pyramidal_4x_SEGMENTATION.tif | 445 | 101.2684221 | 4.394262206 |
| Beta | Saline | P0-4 | WS31925-10C | WS31925_10C_pyramidal_4x_SEGMENTATION.tif | 379 | 94.01090472 | 4.031447215 |
| Beta | Gent | P0-4 | WS31925-11E | WS31925_11E_pyramidal_4x_SEGMENTATION.tif | 387 | 71.67655809 | 5.399254796 |
| Beta | Saline | P0-4 | WS31925-6C | WS31925_6C_pyramidal_4x_SEGMENTATION.tif | 334 | 76.67584219 | 4.35600041 |
| Beta | Gent | P0-4 | WS31925-9E | WS31925_9E_pyramidal_4x_SEGMENTATION.tif | 332 | 64.89282985 | 5.116127633 |

Supplemental Table 3

| Treatment days (P) | Prenatal and postnatal exposures | sample | Min | Max | Mean | StdDev | Median | Percentile 25% | Percentile 75% | No. of Gloms Analyzed |
|---|---|---|---|---|---|---|---|---|---|---|
| 0-4 | none-saline | RC-2C | 995805.6 | 32009522 | 1840846.12 | 703749 | 1729928.5 | 1400832 | 2105344 | 29966 |
| 0-4 | none-saline | RC-4C | 991701.4 | 192889712 | 1701154.87 | 1416238.38 | 1537719.13 | 1245184 | 1900544 | 24750 |
| 0-4 | none-saline | RC-6C | 996093.1 | 6644935 | 1364998.25 | 312983.812 | 1296402.5 | 1140736 | 1501184 | 16738 |
| 0-4 | none-saline | RC-8C | 989593.6 | 171638976 | 1797423 | 3723217 | 1412318.62 | 1245184 | 1769472 | 9501 |
| 0-4 | none-Gent | RC-1E | 983684.1 | 14515802 | 1789735.62 | 810766.812 | 1596858.25 | 1290240 | 2002944 | 12896 |
| 0-4 | none-Gent | RC-5E | 994966.4 | 5946209 | 1513353.87 | 394563.406 | 1446033.62 | 1222656 | 1710080 | 15713 |
| 0-4 | none-Gent | RC-7E | 975466.3 | 312680960 | 1678288.62 | 3757745.5 | 1424045 | 1179648 | 1703936 | 14267 |
| 0-4 | none-Gent | RC-9E | 996922.8 | 12637495 | 1681374.5 | 640445.25 | 1510040.37 | 1273856 | 1863680 | 23530 |
| 0-4 | Beta-saline | TS-2C | 994926.9 | 43303576 | 1697720 | 1300676.37 | 1379513.12 | 1163264 | 1785856 | 9498 |
| 0-4 | Beta-saline | TS-10C | 997198.7 | 14227369 | 2454774.25 | 1029385.75 | 2278120.25 | 1798144 | 2854912 | 22934 |
| 0-4 | Beta-saline | WS-6C | 987828.25 | 30162880 | 1789804.62 | 878309.81 | 1557084.87 | 1269760 | 2007040 | 21153 |
| 0-4 | Beta-saline | WS-10C | 977051.8 | 13563941 | 1723136.62 | 772834.312 | 1523659.5 | 1232896 | 1937408 | 11490 |
| 0-4 | Beta-Gent | TS-1E | 992888.1 | 8657355 | 1569802.12 | 518411.625 | 1443797.12 | 1222656 | 1759232 | 14288 |
| 0-4 | Beta-Gent | TS-7E | 993758.4 | 15593388 | 1818621.37 | 614711.875 | 1728814.5 | 1388544 | 2101248 | 19480 |
| 0-4 | Beta-Gent | TS-13E | 991895.4 | 8912825 | 1670866.12 | 568988.562 | 1547137 | 1275904 | 1890304 | 17210 |
| 6-10 | none-saline | FC-6C | 980112.2 | 19856760 | 1830843.62 | 957311.625 | 1554317.87 | 1236992 | 2072576 | 11700 |
| 6-10 | none-saline | FC-10C | 982032.9 | 139865984 | 2455743.25 | 2791542.75 | 1802019 | 1376256 | 2686976 | 15632 |
| 6-10 | none-saline | FC-14C | 967237.4 | 123522828 | 2343898 | 14784341 | 1491534.5 | 1572864 | 2621440 | 15329 |
| 6-10 | none-Gent | FC-3E | 982500.9 | 81202856 | 2633696.75 | 3049034.5 | 1930192.75 | 1474560 | 2785280 | 8846 |
| 6-10 | none-Gent | FC-5E | 1000039 | 25349072 | 2368188.25 | 1099918.37 | 2116718.25 | 1695744 | 2695168 | 22076 |
| 6-10 | none-Gent | FC-15E | 997883.7 | 25340212 | 2367315 | 1099223.62 | 2115612.5 | 1695744 | 2695168 | 22074 |
| 6-10 | Beta-saline | SS-6C | 987435.6 | 332908736 | 2385112 | 5532374 | 1909874.5 | 1441792 | 2490368 | 17216 |
| 6-10 | Beta-Gent | SS-3E | 988798.6 | 69497048 | 3047314 | 2306634.75 | 2674996 | 1867776 | 3571712 | 7489 |
| 6-10 | Beta-Gent | SS-7E | 978282.9 | 32444400 | 2447933.25 | 1526932.75 | 2183269.5 | 1564672 | 2891776 | 7072 |
| 6-10 | Beta-Gent | SS-9E | 994474.5 | 50034492 | 2955109.25 | 1609206.5 | 2766497.5 | 2048000 | 3424256 | 9850 |
| 6-10 | Beta-Gent | SS-11E | 975457.9 | 244422208 | 2757738.75 | 4653524 | 2448526.25 | 1900544 | 2949120 | 9851 |

Supplemental Table 4

| AKI Analysis Samples (P0-4 and P6-10 Treatments) | | | | | | | |
|---|---|---|---|---|---|---|---|
| **Group** | **Litter** | **# of pups** | **Prenatal+Postnatal Treatment** | **DOB** | **Collected** | **Pup IDs** | **Sex** |
| P0-4 AKI | E81424 | 8 | None + gent | 8/14/2024 | 8/18/2024 | 1E | F |
| | | | None + gent | 8/14/2024 | 8/18/2024 | 5E | M |
| | | | None + gent | 8/14/2024 | 8/18/2024 | 7E | F |
| | | | None + gent | 8/14/2024 | 8/18/2024 | 13E | F |
| | | | None + saline | 8/14/2024 | 8/18/2024 | 4C | F |
| | | | None + saline | 8/14/2024 | 8/18/2024 | 10C | F |
| | | | None + saline | 8/14/2024 | 8/18/2024 | 12C | M |
| | | | None + saline | 8/14/2024 | 8/18/2024 | 16C | M |
| | B5824 | 8 | None + gent | 5/8/2024 | 5/12/2024 | 1E | M |
| | | | None + gent | 5/8/2024 | 5/12/2024 | 3E | M |
| | | | None + gent | 5/8/2024 | 5/12/2024 | 5E | F |
| | | | None + gent | 5/8/2024 | 5/12/2024 | 9E | M |
| | | | None + saline | 5/8/2024 | 5/12/2024 | 2C | M |
| | | | None + saline | 5/8/2024 | 5/12/2024 | 4C | F |
| | | | None + saline | 5/8/2024 | 5/12/2024 | 6C | F |
| | | | None + saline | 5/8/2024 | 5/12/2024 | 8C | M |
| | N110224 | 12 | None + gent | 11/2/2024 | 11/6/2024 | 1E | M |
| | | | None + gent | 11/2/2024 | 11/6/2024 | 3E | F |
| | | | None + gent | 11/2/2024 | 11/6/2024 | 5E | F |
| | | | None + gent | 11/2/2024 | 11/6/2024 | 7E | M |
| | | | None + gent | 11/2/2024 | 11/6/2024 | 9E | M |
| | | | None + gent | 11/2/2024 | 11/6/2024 | 11E | F |
| | | | None + saline | 11/2/2024 | 11/6/2024 | 2C | F |
| | | | None + saline | 11/2/2024 | 11/6/2024 | 4C | F |
| | | | None + saline | 11/2/2024 | 11/6/2024 | 6C | F |
| | | | None + saline | 11/2/2024 | 11/6/2024 | 8C | F |
| | | | None + saline | 11/2/2024 | 11/6/2024 | 10C | M |
| | | | None + saline | 11/2/2024 | 11/6/2024 | 12C | M |
| | VC31925 | 6 | Saline + gent | 3/19/2025 | 3/23/2025 | 1E | F |
| | | | Saline + gent | 3/19/2025 | 3/23/2025 | 3E | M |
| | | | Saline + gent | 3/19/2025 | 3/23/2025 | 5E | M |
| | | | Saline + gent | 3/19/2025 | 3/23/2025 | 7E | M |
| | | | Saline + saline | 3/19/2025 | 3/23/2025 | 4C | F |
| | | | Saline + saline | 3/19/2025 | 3/23/2025 | 6C | F |
| | DS81424 | 8 | Beta + gent | 8/14/2024 | 8/18/2024 | 1E | F |
| | | | Beta + gent | 8/14/2024 | 8/18/2024 | 5E | M |
| | | | Beta + gent | 8/14/2024 | 8/18/2024 | 7E | F |
| | | | Beta + gent | 8/14/2024 | 8/18/2024 | 13E | F |
| | | | Beta + saline | 8/14/2024 | 8/18/2024 | 4C | F |
| | | | Beta + saline | 8/14/2024 | 8/18/2024 | 10C | F |

| | | | | | | | |
|---|---|---|---|---|---|---|---|
| | | | Beta + saline | 8/14/2024 | 8/18/2024 | 12C | M |
| | | | Beta + saline | 8/14/2024 | 8/18/2024 | 16C | M |
| | CS71724 | 5 | Beta + gent | 7/17/2024 | 7/21/2024 | 1E | F |
| | | | Beta + gent | 7/17/2024 | 7/21/2024 | 3E | M |
| | | | Beta + saline | 7/17/2024 | 7/21/2024 | 2C | F |
| | | | Beta + saline | 7/17/2024 | 7/21/2024 | 4C | M |
| | | | Beta + saline | 7/17/2024 | 7/21/2024 | 6C | F |
| | WS31925 | 5 | Beta + gent | 3/19/2025 | 3/23/2025 | 1E | M |
| | | | Beta + gent | 3/19/2025 | 3/23/2025 | 5E | M |
| | | | Beta + gent | 3/19/2025 | 3/23/2025 | 7E | F |
| | | | Beta + saline | 3/19/2025 | 3/23/2025 | 4C | F |
| | | | Beta + saline | 3/19/2025 | 3/23/2025 | 8C | F |
| P0-4 AKI (no postnat al) | AI-C91725 | 2 | Saline + NA | 9/17/2025 | 9/21/2025 | 2C | M |
| | | | Saline + NA | 9/17/2025 | 9/21/2025 | 4C | M |
| | AH-S91725 | 2 | Saline + NA | 9/17/2025 | 9/21/2025 | 2C | M |
| | | | Saline + NA | 9/17/2025 | 9/21/2025 | 4C | F |
| P6-10 AKI | F91224 | 7 | None + gent | 9/12/2024 | 9/22/2024 | 1E | M |
| | | | None + gent | 9/12/2024 | 9/22/2024 | 7E | F |
| | | | None + gent | 9/12/2024 | 9/22/2024 | 13E | M |
| | | | None + saline | 9/12/2024 | 9/22/2024 | 2C | F |
| | | | None + saline | 9/12/2024 | 9/22/2024 | 4C | F |
| | | | None + saline | 9/12/2024 | 9/22/2024 | 8C | M |
| | | | None + saline | 9/12/2024 | 9/22/2024 | 12C | M |
| | IC92524 | 9 | None + gent | 9/5/2024 | 10/5/2024 | 1E | F |
| | | | None + gent | 9/5/2024 | 10/5/2024 | 3E | M |
| | | | None + gent | 9/5/2024 | 10/5/2024 | 7E | F |
| | | | None + gent | 9/5/2024 | 10/5/2024 | 11E | M |
| | | | None + gent | 9/5/2024 | 10/5/2024 | 17E | M |
| | | | None + saline | 9/5/2024 | 10/5/2024 | 10C | F |
| | | | None + saline | 9/5/2024 | 10/5/2024 | 12C | M |
| | | | None + saline | 9/5/2024 | 10/5/2024 | 14C | M |
| | | | None + saline | 9/5/2024 | 10/5/2024 | 16C | M |
| | QC122524 | 6 | Saline + gent | 12/25/2024 | 1/4/2025 | 1E | M |
| | | | Saline + gent | 12/25/2024 | 1/4/2025 | 3E | M |
| | | | Saline + gent | 12/25/2024 | 1/4/2025 | 9E | F |
| | | | Saline + saline | 12/25/2024 | 1/4/2025 | 2C | M |
| | | | Saline + saline | 12/25/2024 | 1/4/2025 | 4C | M |
| | | | Saline + saline | 12/25/2024 | 1/4/2025 | 10C | F |
| | M102324 | 15 | None + gent | 10/23/2024 | 11/2/2024 | 1E | M |
| | | | None + gent | 10/23/2024 | 11/2/2024 | 3E | M |
| | | | None + gent | 10/23/2024 | 11/2/2024 | 5E | M |
| | | | None + gent | 10/23/2024 | 11/2/2024 | 7E | M |
| | | | None + gent | 10/23/2024 | 11/2/2024 | 9E | F |
| | | | None + gent | 10/23/2024 | 11/2/2024 | 11E | F |

| | | | | | | | |
|---|---|---|---|---|---|---|---|
| | | | None + gent | 10/23/2024 | 11/2/2024 | 13E | F |
| | | | None + gent | 10/23/2024 | 11/2/2024 | 15E | F |
| | | | None + saline | 10/23/2024 | 11/2/2024 | 2C | F |
| | | | None + saline | 10/23/2024 | 11/2/2024 | 4C | F |
| | | | None + saline | 10/23/2024 | 11/2/2024 | 6C | F |
| | | | None + saline | 10/23/2024 | 11/2/2024 | 8C | F |
| | | | None + saline | 10/23/2024 | 11/2/2024 | 10C | M |
| | | | None + saline | 10/23/2024 | 11/2/2024 | 12C | M |
| | | | None + saline | 10/23/2024 | 11/2/2024 | 14C | M |
| | CS71724 | 3 | Beta + gent | 7/17/2024 | 7/27/2024 | 9E | M |
| | | | Beta + gent | 7/17/2024 | 7/27/2024 | 11E | F |
| | | | Beta + saline | 7/17/2024 | 7/27/2024 | 8C | M |
| | JS92524 | 8 | Beta + gent | 9/25/2024 | 10/5/2024 | 5E | M |
| | | | Beta + gent | 9/25/2024 | 10/5/2024 | 9E | F |
| | | | Beta + gent | 9/25/2024 | 10/5/2024 | 11E | M |
| | | | Beta + gent | 9/25/2024 | 10/5/2024 | 13E | M |
| | | | Beta + saline | 9/25/2024 | 10/5/2024 | 2C | F |
| | | | Beta + saline | 9/25/2024 | 10/5/2024 | 6C | F |
| | | | Beta + saline | 9/25/2024 | 10/5/2024 | 8C | M |
| | | | Beta + saline | 9/25/2024 | 10/5/2024 | 16C | M |
| | PS122524 | 6 | Beta + gent | 12/25/2024 | 1/4/2025 | 1E | M |
| | | | Beta + gent | 12/25/2024 | 1/4/2025 | 3E | M |
| | | | Beta + gent | 12/25/2024 | 1/4/2025 | 9E | F |
| | | | Beta + saline | 12/25/2024 | 1/4/2025 | 2C | M |
| | | | Beta + saline | 12/25/2024 | 1/4/2025 | 8C | F |
| | | | Beta + saline | 12/25/2024 | 1/4/2025 | 10C | F |

**Adult AKI samples**

| Group | Litter | # of rats | Treatment | DOB | Collected | Rat IDs | Sex |
|---|---|---|---|---|---|---|---|
| Adult AKI | Adult Breeders | 8 | x5 days gent | | 10/29/2025 | Breeder#1M | M |
| | | | x5 days gent | | 10/29/2025 | Breeder#3M | M |
| | | | x5 days gent | | 10/29/2025 | Breeder#8F | F |
| | | | x5 days gent | | 10/29/2025 | Breeder#10F | F |
| | | | x5 days saline | | 10/29/2025 | Breeder#2M | M |
| | | | x5 days saline | | 10/29/2025 | Breeder#4M | M |
| | | | x5 days saline | | 10/29/2025 | Breeder#4F | F |
| | | | x5 days saline | | 10/29/2025 | Breeder#6F | F |

**2 Week Analysis Samples (P0-4 and P6-10 Treatments)**

| Group | Litter | # of pups | Prenatal+Postnatal Treatment | DOB | Collected | Pup IDs | Sex |
|---|---|---|---|---|---|---|---|
| P0-4 2weeks | E81424 | 6 | None + gent | 8/14/2024 | 9/1/2024 | 1E | M |
| | | | None + gent | 8/14/2024 | 9/1/2024 | 3E | M |
| | | | None + gent | 8/14/2024 | 9/1/2024 | 13E | M |
| | | | None + saline | 8/14/2024 | 9/1/2024 | 6C | M |
| | | | None + saline | 8/14/2024 | 9/1/2024 | 8C | M |
| | | | None + saline | 8/14/2024 | 9/1/2024 | 10C | M |
| | B5824 | 4 | None + gent | 5/8/2024 | 5/26/2024 | 7E | F |
| | | | None + gent | 5/8/2024 | 5/26/2024 | 11E | F |
| | | | None + saline | 5/8/2024 | 5/26/2024 | 10C | M |
| | | | None + saline | 5/8/2024 | 5/26/2024 | 12C | M |
| | UC21825 | 11 | Saline + gent | 2/18/2025 | 3/8/2025 | 1E | M |
| | | | Saline + gent | 2/18/2025 | 3/8/2025 | 3E | M |
| | | | Saline + gent | 2/18/2025 | 3/8/2025 | 5E | F |
| | | | Saline + gent | 2/18/2025 | 3/8/2025 | 7E | F |
| | | | Saline + gent | 2/18/2025 | 3/8/2025 | 9E | F |
| | | | Saline + gent | 2/18/2025 | 3/8/2025 | 11E | M |
| | | | Saline + saline | 2/18/2025 | 3/8/2025 | 2C | F |
| | | | Saline + saline | 2/18/2025 | 3/8/2025 | 4C | F |
| | | | Saline + saline | 2/18/2025 | 3/8/2025 | 6C | M |
| | | | Saline + saline | 2/18/2025 | 3/8/2025 | 8C | M |
| | | | Saline + saline | 2/18/2025 | 3/8/2025 | 10C | M |
| | VC31925 | 1 | Saline + saline | 3/19/2025 | 4/6/2025 | 2C | F |
| | DS81424 | 7 | Saline + gent | 8/14/2024 | 9/1/2024 | 3E | M |
| | | | Saline + gent | 8/14/2024 | 9/1/2024 | 9E | F |
| | | | Saline + gent | 8/14/2024 | 9/1/2024 | 15E | F |
| | | | Saline + saline | 8/14/2024 | 9/1/2024 | 2C | M |
| | | | Saline + saline | 8/14/2024 | 9/1/2024 | 6C | M |
| | | | Saline + saline | 8/14/2024 | 9/1/2024 | 8C | M |
| | | | Saline + saline | 8/14/2024 | 9/1/2024 | 14C | M |
| | TS21825 | 6 | Saline + gent | 2/18/2025 | 3/8/2025 | 3E | F |
| | | | Saline + gent | 2/18/2025 | 3/8/2025 | 5E | F |
| | | | Saline + gent | 2/18/2025 | 3/8/2025 | 11E | F |
| | | | Saline + gent | 2/18/2025 | 3/8/2025 | 15E | M |
| | | | Saline + saline | 2/18/2025 | 3/8/2025 | 6C | F |
| | | | Saline + saline | 2/18/2025 | 3/8/2025 | 8C | F |
| | WS31925 | 2 | Saline + gent | 3/19/2025 | 4/6/2025 | 3E | M |
| | | | Saline + saline | 3/19/2025 | 4/6/2025 | 2C | F |
| P6-10 2weeks | IC92524 | 8 | Saline + gent | 9/25/2024 | 10/5/2024 | 5E | M |
| | | | Saline + gent | 9/25/2024 | 10/5/2024 | 9E | F |
| | | | Saline + gent | 9/25/2024 | 10/5/2024 | 13E | M |
| | | | Saline + gent | 9/25/2024 | 10/5/2024 | 15E | M |
| | | | Saline + saline | 9/25/2024 | 10/5/2024 | 2C | F |

| | | | | | | | |
|---|---|---|---|---|---|---|---|
| | | | Saline + saline | 9/25/2024 | 10/5/2024 | 4C | F |
| | | | Saline + saline | 9/25/2024 | 10/5/2024 | 6C | F |
| | | | Saline + saline | 9/25/2024 | 10/5/2024 | 8C | M |
| | | | Saline + saline | 9/25/2024 | 10/5/2024 | 12C | M |
| | QC122524 | 6 | Saline + gent | 12/25/2024 | 1/18/2025 | 5E | M |
| | | | Saline + gent | 12/25/2024 | 1/18/2025 | 7E | M |
| | | | Saline + saline | 12/25/2024 | 1/18/2025 | 6C | M |
| | | | Saline + saline | 12/25/2024 | 1/18/2025 | 8C | M |
| | | | Saline + saline | 12/25/2024 | 1/18/2025 | 12C | F |
| | VC31925 | 2 | Saline + gent | 3/19/2025 | 4/12/2025 | 11E | F |
| | | | Saline + gent | 3/19/2025 | 4/12/2025 | 13E | F |
| | JS90524 | 8 | Beta + gent | 9/5/2024 | 10/19/2024 | 1E | F |
| | | | Beta + gent | 9/5/2024 | 10/19/2024 | 3E | M |
| | | | Beta + gent | 9/5/2024 | 10/19/2024 | 7E | F |
| | | | Beta + gent | 9/5/2024 | 10/19/2024 | 15E | F |
| | | | Beta + saline | 9/5/2024 | 10/19/2024 | 4C | F |
| | | | Beta + saline | 9/5/2024 | 10/19/2024 | 10C | F |
| | | | Beta + saline | 9/5/2024 | 10/19/2024 | 12C | M |
| | | | Beta + saline | 9/5/2024 | 10/19/2024 | 14C | M |
| | PS122524 | 6 | Beta + gent | 12/25/2024 | 1/18/2025 | 7E | M |
| | | | Beta + gent | 12/25/2024 | 1/18/2025 | 11E | F |
| | | | Beta + saline | 12/25/2024 | 1/18/2025 | 4C | M |
| | | | Beta + saline | 12/25/2024 | 1/18/2025 | 6C | M |
| | | | Beta + saline | 12/25/2024 | 1/18/2025 | 12C | F |
| | | | Beta + saline | 12/25/2024 | 1/18/2025 | 14C | F |
| | SS20525 | 3 | Beta + gent | 2/5/2025 | 3/1/2025 | 5E | M |
| | | | Beta + gent | 2/5/2025 | 3/1/2025 | 1E | F |
| | | | Beta + saline | 2/5/2025 | 3/1/2025 | 2C | F |
| | WS31925 | 1 | Beta + gent | 3/19/2025 | 4/12/2025 | 13E | F |

**Adult Analysis Samples (P0-4 and P6-10 Treatments)**

| Group | Litter | # of pups | Prenatal+Postnatal Treatment | DOB | Collected | Pup IDs | Sex |
|---|---|---|---|---|---|---|---|
| P0-4 3mo | R11025 | 10 | None + gent | 1/10/2025 | 4/10/2025 | 1E | M |
| | | | None + gent | 1/10/2025 | 4/10/2025 | 3E | M |
| | | | None + gent | 1/10/2025 | 4/10/2025 | 5E | M |
| | | | None + gent | 1/10/2025 | 4/10/2025 | 7E | F |
| | | | None + gent | 1/10/2025 | 4/10/2025 | 9E | F |
| | | | None + saline | 1/10/2025 | 4/10/2025 | 2C | M |
| | | | None + saline | 1/10/2025 | 4/10/2025 | 4C | M |
| | | | None + saline | 1/10/2025 | 4/10/2025 | 6C | M |
| | | | None + saline | 1/10/2025 | 4/10/2025 | 8C | F |
| | | | None + saline | 1/10/2025 | 4/10/2025 | 10C | F |
| | VC31925 | 2 | Saline + gent | 3/19/2025 | 6/19/2025 | 8C | M |
| | | | Saline + saline | 3/19/2025 | 6/19/2025 | 9E | M |

| | | | | | | | |
|---|---|---|---|---|---|---|---|
| | TS21825 | 8 | Saline + gent | 2/18/2025 | 5/18/2025 | 1E | F |
| | | | Saline + gent | 2/18/2025 | 5/18/2025 | 7E | M |
| | | | Saline + gent | 2/18/2025 | 5/18/2025 | 9E | M |
| | | | Saline + gent | 2/18/2025 | 5/18/2025 | 13E | M |
| | | | Saline + saline | 2/18/2025 | 5/18/2025 | 2C | F |
| | | | Saline + saline | 2/18/2025 | 5/18/2025 | 4C | M |
| | | | Saline + saline | 2/18/2025 | 5/18/2025 | 10C | M |
| | | | Saline + saline | 2/18/2025 | 5/18/2025 | 12C | F |
| | WS31925 | 4 | Beta + gent | 3/19/2025 | 6/19/2025 | 9E | F |
| | | | Beta + gent | 3/19/2025 | 6/19/2025 | 11E | F |
| | | | Beta + saline | 3/19/2025 | 6/19/2025 | 6C | M |
| | | | Beta + saline | 3/19/2025 | 6/19/2025 | 10C | F |
| P6-10 3mo | SS20525 | 8 | Beta + gent | 2/5/2025 | 5/15/2025 | 3E | F |
| | | | Beta + gent | 2/5/2025 | 5/15/2025 | 7E | M |
| | | | Beta + gent | 2/5/2025 | 5/15/2025 | 9E | M |
| | | | Beta + gent | 2/5/2025 | 5/15/2025 | 11E | M |
| | | | Beta + saline | 2/5/2025 | 5/15/2025 | 4C | F |
| | | | Beta + saline | 2/5/2025 | 5/15/2025 | 6C | M |
| | | | Beta + saline | 2/5/2025 | 5/15/2025 | 8C | M |
| | | | Beta + saline | 2/5/2025 | 5/15/2025 | 10C | M |
| | F91224 | 8 | None + gent | 9/12/2024 | 12/12/2024 | 3E | F |
| | | | None + gent | 9/12/2024 | 12/12/2024 | 5E | M |
| | | | None + gent | 9/12/2024 | 12/12/2024 | 9E | M |
| | | | None + gent | 9/12/2024 | 12/12/2024 | 15E | F |
| | | | None + saline | 9/12/2024 | 12/12/2024 | 6C | F |
| | | | None + saline | 9/12/2024 | 12/12/2024 | 10C | M |
| | | | None + saline | 9/12/2024 | 12/12/2024 | 14C | M |
| | | | None + saline | 9/12/2024 | 12/12/2024 | 16C | F |

Supplemental Table 5

| Primer | Sequence |
|---|---|
| Gapdh-a forward | 5'-GACAGCCGCATCTTCTTGTG |
| Gapdh-a reverse | 5'-GGTAACCAGGCGTCCGATAC |
| Gapdh-b forward | 5'-GACAGTCAGCCGCATCTTCT |
| Gapdh-b reverse | 5'-TTAAAAGCAGCCCTGGTGAC |
| Ppia-b forward | 5'-CCAAACACAAATGGTTCCCAGT |
| Ppia-b reverse | 5'-ATTCCTGGACCCAAAACGCT |
| Kim1-a forward | 5'-GTCGTTGTGATTCCTCCACG |
| Kim1-a reverse | 5'-TGGCTCTATGTGAGCAAGGAC |
| Sry forward | 5'-AAGCGCCCCSTGAATGCAT |
| Sry reverse | 5'-CGATGAGGCTGATATTTATA |

Supplemental Table 6: *RT-qPCR For P4 Rat Sex Determination*

| Sample Name | Ave CT/ppia-b | Ave CT/SRY | Sex |
|---|---|---|---|
| VC-4C | 15.623 | NA | Female |
| VC-6C | 15.197 | NA | Female |
| VC-1E | 29.539 | 30.536 | Male |
| VC-3E | 15.032 | 30.079 | Male |
| VC-5E | 14.784 | 30.544 | Male |
| VC-7E | 14.571 | 28.878 | Male |
| WS-4C | 14.231 | NA | Female |
| WS-8C | 14.899 | NA | Female |
| WS-1E | 13.646 | 28.387 | Male |
| WS-5E | 16.636 | NA | Female |
| WS-7E | 15.112 | NA | Female |
| Con81424-2C | 16.582 | 31.265 | Male |
| Con81424-4C | 15.713 | NA | Female |
| Con81424-12C | 14.706 | 27.429 | Male |
| Con81424-5G | 13.964 | 26.578 | Male |
| Con81424-7G | 15.112 | 30.765 | Male |
| Con81424-9G | 14.886 | 29.296 | Male |
| Con81424-11G | 14.921 | 29.357 | Male |
| Ste81424-4C | 33.134 | NA | Female |
| Ste81424-10C | 14.142 | NA | Female |
| Ste81424-12C | 15.058 | 27.531 | Male |
| Ste81424-16C | 15.068 | 27.763 | Male |
| Ste81424-1G | 13.818 | 27.901 | Male |
| Ste81424-5G | 30.887 | NA | Female |
| Ste81424-7G | 14.884 | NA | Female |
| Ste81424-13G | 15.895 | 34.280 | Male |
| ET5824-1G | 15.571 | 26.619 | Male |
| ET5824-2C | 14.420 | 27.807 | Male |
| ET5824-3G | 14.779 | 26.164 | Male |
| ET5824-4C | 15.086 | NA | Female |
| ET5824-5G | 15.590 | NA | Female |
| ET5824-6C | 15.095 | NA | Female |
| ET5824-9G | 14.207 | 25.683 | Male |
| ET5824-8C | 15.422 | NA | Female |
| AI-C91725-2C | 16.333 | 33.405 | Male |
| AI-C91725-4C | 17.741 | 29.694 | Male |
| AH-S91725-2C | 17.268 | 33.246 | Male |
| AH-S91725-4C | 15.439 | NA | Female |
| Adult-2MC | 22.354 | 28.255 | Male |
| Adult-4FC | 21.386 | NA | Female |
| Adult-8MC | 17.447 | 31.654 | Male |
| Adult-12FC | 17.615 | NA | Female |

**Supplemental Movie 1**. Three-dimensional rendering of a light-sheet microscopy image of cleared control kidney with lectin-labeled glomeruli.
